# First report of genetic transformation and CRISPR/Cas12a-mediated gene editing of European beech (*Fagus sylvatica* L.) employing a transient protoplast system

**DOI:** 10.64898/2026.02.06.703565

**Authors:** Virginia Zahn, Alice-Jeannine Sievers, Birgit Kersten, Matthias Fladung, Tobias Bruegmann

## Abstract

*Fagus sylvatica* L. (European beech) is a dominant hardwood forest tree species across Central Europe, supporting diverse ecosystem services and forming the basis of a significant market for high-value timber. However, climate change increasingly threatens beech vitality and productivity, making molecular insights into its stress resilience and functional validation of underlying genes urgently needed. Here, we report a protocol for protoplast isolation from seedling leaves and demonstrate, for the first time, transient genetic transformation and CRISPR/Cas-mediated genome editing in *F. sylvatica*. PEG-mediated transformation was sequentially optimized, achieving efficiencies of 59 ± 6.19% within distinct seasonal windows. Protoplast yield and transformation efficiency showed pronounced temporal variation throughout the year, indicating a strong seasonal influence on reproducibility of the workflow despite controlled growth conditions. A basic molecular toolkit for functional genomics and future biotechnological applications was established by testing a set of promoters and reporters. For proof-of-concept genome editing, we achieved 4.75 to 32.69% editing efficiencies in the *PHYTOENE DESATURASE* gene (*FsPDS*) using temperature-tolerant *Lb*Cas12a (tt*Lb*Cas12a). Although further optimization of transformation reproducibility and regeneration systems remains necessary, the presented protoplast platform provides a valuable foundation for transient functional assays and genome editing studies in this non-model tree species.

## Introduction

European beech (*Fagus sylvatica* L.), beech for short, is one of the most widespread deciduous tree species in Central Europe, forming stable climax stands across broad latitudinal and longitudinal gradients (Fuchs et al., 2024; Houston Durrant et al., 2016). Combining ecological and economic values, beech has long been considered a cornerstone of near-natural, climate-resilient forestry in Central Europe because of its phenotypic plasticity and broad ecological amplitude (Felton et al., 2010; Wühlisch, 2008). However, beech is increasingly threatened by climate change, as evidenced by widespread crown defoliation (Bussotti et al., 2024; Corcobado et al., 2020; Langer and Bußkamp, 2023), with 82% of monitored beech trees exhibiting at least moderate crown defoliation symptoms in the 2024 German forest condition survey (BMLEH [Bundesministerium für Landwirtschaft, Ernährung und Heimat], 2025). In Germany alone, the economic loss caused by the decline of beech trees is projected to range from €180 billion up to €4 trillion, depending on the scenario (Baumbach et al., 2019). Understanding the molecular basis of climate resilience in beech has thus become an urgent research priority (Pfenninger et al., 2021).

Forest trees lag far behind annual crops in the application of molecular tools (Borthakur et al., 2022). Breeding programs for forest trees still rely largely on phenotypic selection, and gene function studies via transgenic approaches are nascent even in the model tree genus *Populus* (Sulis et al., 2023). This gap is compounded by long generation times (approx. 50 years in beech), large heterozygous genomes lacking robust reference assemblies, and recalcitrance to *in vitro* regeneration (Houston Durrant et al., 2016; Wang et al., 2025). Although a high-quality reference genome for beech has recently become available, revealing a diploid genome comprising 12 chromosomes and a total assembly size of approx. 541 Mb, its gene models still lack the depth of experimental validation (Mishra et al., 2021).

Protoplasts, single plant cells whose cell wall has been enzymatically degraded, enable rapid, regeneration-independent functional assays at the cell level (Yoo et al., 2007). PEG-mediated protoplast transformation is the most widely used method for DNA uptake, allowing direct testing of gene constructs, including promoter-reporter assays or constructs encoding Cas nucleases and guide-RNAs for genome editing (Panda et al., 2024). This approach is especially valuable for beech, whose high recalcitrance to *in vitro* culture and whole-plant regeneration has been recognized since the late 1970s and continues to engage current research (Chalupa, 1979; Thiesen et al., 2025; Zahn et al., 2025). When combined with plant regeneration, protoplast-based systems enable DNA-free genome editing via ribonucleoprotein (RNP) delivery (Marques et al., 2025). This approach is particularly attractive for long-lived tree species such as beech because it avoids stable integration of transgenes and circumvents lengthy breeding or segregation steps, thereby simplifying subsequent breeding and regulatory processes (Tuncel et al., 2025). Earlier protocols were limited to the isolation of protoplasts from freshly emerged leaves of field-grown trees (Ahuja, 1984; Lang and Kohlenbach, 1988). To date, protoplast transformation has not been reported for beech, despite its efficient application in many other forest tree species (Lu et al., 2025; Pavese et al., 2024).

CRISPR/Cas-mediated gene editing has revolutionized functional genomics across diverse plant species (Jinek et al., 2012; Zetsche et al., 2015). Among forest trees, the use of CRISPR/Cas12a was only reported in initial studies in poplar (An et al., 2020; Das et al., 2025). Compared to the commonly used Cas9, Cas12a offers advantages such as recognizing a T-rich protospacer-adjacent motif (PAM) sequence, producing staggered DNA cuts and larger deletions. The compact target-determining CRISPR-RNA (crRNA) architecture and smaller overall construct size are advantageous for transformation systems with probably limited efficiency. In addition, the shorter crRNA size and the ability of Cas12a to process multiplexed crRNA arrays facilitate the simultaneous targeting of multiple genes, which is particularly relevant for tree species characterized by genomic heterogeneity and extensive gene duplications as well as for the engineering of complex polygenic traits (Zetsche et al., 2015). Another advantage over Cas9 and the standard Cas12a lies in the temperature-tolerant tt*Lb*Cas12a variant that we employed, which enhances editing efficiency at the lower temperatures typical of plant tissue culture (Malzahn et al., 2019; Schindele and Puchta, 2020). Multiple crRNAs were selected based on predicted secondary structure to maximize editing efficiency (Creutzburg et al., 2020; Zhu and Liang, 2019). The *PHYTOENE DESATURASE* gene (*PDS*) was identified in beech and subsequently used as the target gene. *PDS* encodes a key enzyme in the carotenoid biosynthesis pathway that is frequently used as a model editing locus due to its potential for visual phenotypes in regenerated plants and ease of identification due to high sequence conservation (Fan et al., 2015; Hirschberg, 2001; Pavese et al., 2021).

Here, we first present a protocol for efficient protoplast isolation from beech seedling leaves with high yield and vitality. We then report the first successful transient transformation in beech, systematically optimizing key parameters of PEG-mediated DNA uptake. As a step towards building a molecular toolbox for beech, we successfully demonstrated the functionality of a panel of constitutive promoters driving the expression of fluorescent proteins. Finally, using tt*Lb*Cas12a and four crRNAs, we achieved targeted nucleotide deletions in the *FsPDS* gene, marking the first successful transient genome editing in European beech.

## Results

### Protoplast isolation and cultivation

Across the tested time periods, leaves collected during winter to late spring (January–May) and autumn (September–November) were most suitable for protoplast isolation. Using the final protocol, an average yield of 5.5 × 10T protoplasts g⁻¹ FW (SD = 3.6 × 10T) with a mean viability of 94.2% (SD = 3.5%) was obtained, with maximum values reaching 1.5 × 108 protoplasts g⁻¹ FW (Figure 1A; Table S1). A subset of protoplasts exhibited strong viability fluorescence despite lacking chloroplasts, and rare dividing protoplasts were observed. Calcofluor White staining confirmed complete cell wall removal following isolation (Figure 1B). Cell wall regeneration initiated within 1 dpi, with uniform fluorescence indicating full reformation by 7 dpi. The cells subsequently proliferated, forming dense aggregates and microcolonies by 12 weeks (Figure 1C). However, further development beyond the microcallus stage was not observed under the applied culture conditions. Contamination occurred in 5 of 126 wells (3.97%) and was restricted to two of twelve isolations (Table S1). In initial tests, antibiotics did not reduce cell viability, whereas addition of the fungicide natamycin caused extensive cell death across multiple replicates (Figure S1).

**Figure 1:**
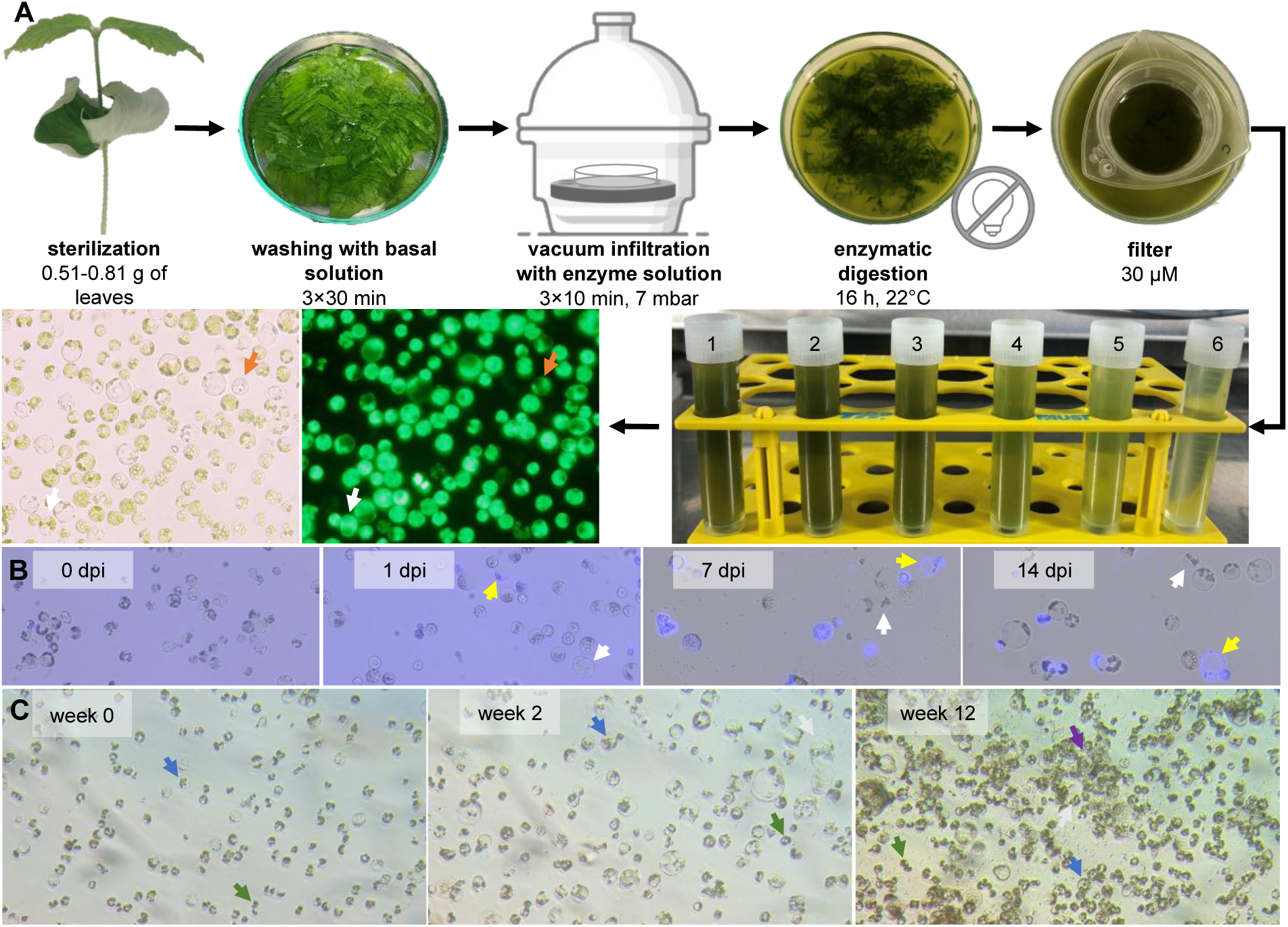
Isolation and cultivation of protoplasts derived from beech seedling leaves. (A) Seedlings were grown *ex vitro*. The primary leaves were harvested, surface-sterilized, cut into strips, and washed in basal solution. Subsequently, tissue was vacuum-infiltrated with enzyme solution and digested overnight. The resulting suspension was filtered (tubes 1-3), and residual tissue was washed three times with W5 solution to release additional protoplasts (tubes 4-6). Filtrates were centrifuged, and the pooled pellets were resuspended in W5 solution. The bright-field image (left) shows a heterogeneous population consisting of chloroplast-containing and chloroplast-free protoplasts (orange arrow). The FDA staining (right) confirmed the high viability of all observed protoplasts, as indicated by strong green fluorescence. The white arrow highlights the rare observation of a dividing protoplast detected within the isolate at day 0. (B) Calcofluor White staining confirming remaining cell wall fragments by absence of fluorescence in freshly isolated protoplasts (0 dpi). Within one day, the first signs of wall regeneration (yellow arrow) and occasional cell divisions (white arrow) were observed. By 7 dpi, cells displayed uniform blue fluorescence, indicating full cell wall regeneration. Staining at 14 dpi confirmed the continued presence of the cell wall. (C) Changes in protoplast morphology and growth over time. Representative images at day 0, week 2, and week 12 in cultivation medium. At day 0, all viable protoplasts are spherical single cells (blue arrow). By week 2, cells have begun to divide (white arrow). By week 12, multicellular clusters (purple arrow) have formed. Over time, the presence of dead cells and cellular debris (green arrow) increases.

### Promoter-reporter validation

To establish a fundamental molecular toolbox for beech, a set of promoter-reporter constructs was tested via transient expression in isolated protoplasts. This marks the first successful molecular characterization of regulatory elements in this species. Successful transient expression of *mEGFP* or *turboRFP* was achieved using six different constitutive promoters (Figure 2A–F).

**Figure 2:**
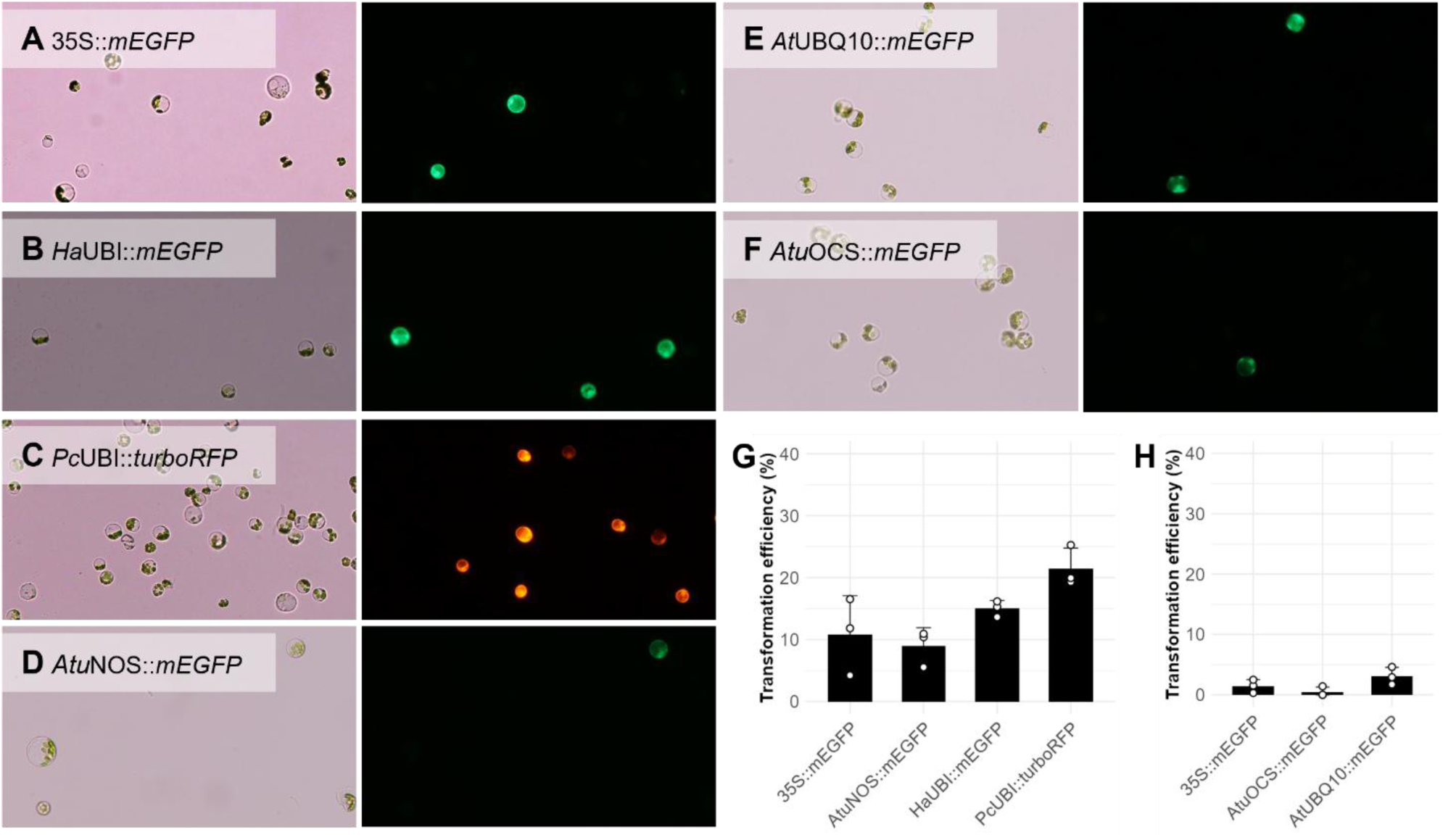
Evaluation of reporter constructs in beech protoplasts. (A–F) Bright-field (left) and fluorescence (right) images of protoplasts 24 h after PEG-mediated transformation. (A–D) Transformations using parameter set A (17/03/2025) with 35S::*mEGFP*, *Ha*UBI::*mEGFP*, *Atu*NOS::*mEGFP*, and *Pc*UBI::*turboRFP*. (E, F) Transformations using parameter set G (24/10/2025) with *At*UBQ10::*mEGFP* and *Atu*OCS::*mEGFP*. No autofluorescence was detected in untransformed protoplasts under identical imaging conditions (Figure S2). (G, H) Transformation efficiencies of vectors transformed on (G) 17/03/2025 and (H) 24/10/2025. Dots represent technical replicates (n = 3), bars indicate mean transformation efficiency, and error bars represent SD (Table S2).

The tested constructs yielded markedly different frequencies of detectable fluorescent cells (transformation efficiency) across two independent transformation days (Figure 2G, H). The *Pc*UBI promoter, driving *turboRFP* expression, achieved the highest overall transformation efficiency observed in this study, exceeding 20% (Figure 2G). All three tested *UBIQUITIN* promoters (*Ha*UBI, *Pc*UBI, and *At*UBQ10) generally resulted in higher frequencies of fluorescent cells than the CaMV35S promoter. Conversely, the *A. tumefaciens*-derived promoters (*Atu*NOS and *Atu*OCS) yielded lower transformation efficiencies than the *CaMV*35S promoter on both transformation days. Both mEGFP and turboRFP reporter proteins were successfully expressed and fluorescently detectable, confirming their functionality in beech cells.

### PEG-mediated transformation and optimization

Seven independent transformation experiments (I–VII) were conducted to systematically optimize the PEG-mediated protocol by testing individual parameters stepwise while carrying forward the best-performing conditions from previous experiments (Figure 3; Table 1). Parameter sets A–R are summarized in Figure 3, whereas additional PEG concentration conditions (S–V) are reported separately in Table 1 for clarity and to maintain readability of the workflow overview. Pre-incubation of protoplasts with plasmid DNA on ice was found to be critical, with extending the pre-incubation to 30 min yielding the highest efficiency (Figure 3A I, VI). Optimal cell density was determined to be 2 × 10ª cells/mL using a total amount of 5 × 10r protoplasts, which outperformed the lower density conditions. At this optimal density, increasing the DNA (25 µg vs 50 µg) or protoplast amount (1 × 10^6^) resulted in minor reductions in efficiency (Figure 3A II). Further increasing the cell density by reducing the transfection buffer volume decreased efficiency (Figure 3A VIII). MMG buffer resulted in higher transformation rates than W5 solution, irrespective of the PEG type used (Figure 3A III). Regarding the PEG treatment, PEG_1500_ slightly outperformed PEG_4000_ (Figure 3A III, IV). Within the PEG types, 5 min and 10 min incubations yielded similar high efficiencies, while increasing the incubation time to 15 min led to reduced efficiencies (Figure 3III). However, when the DNA pre-incubation on ice was extended to 30 min the 5 min PEG incubation produced higher transformation efficiencies than the 10 min incubation (Figure 3A VI). Additional parameter sets not included in the sequential workflow overview (S–V) further demonstrated that reducing the PEG concentration from 40% to 30% or 20% decreased transformation efficiencies under all tested conditions. Microscopic inspection confirmed that the decrease in efficiency was not primarily due to morphological damage, as protoplasts largely retained their integrity even after extended 30% PEG treatment (Figure S3). Applying a 42°C heat shock for 1 min immediately after PEG addition negatively impacted transformation efficiency, regardless of subsequent incubation conditions (Figure 3A V).

**Figure 3:**
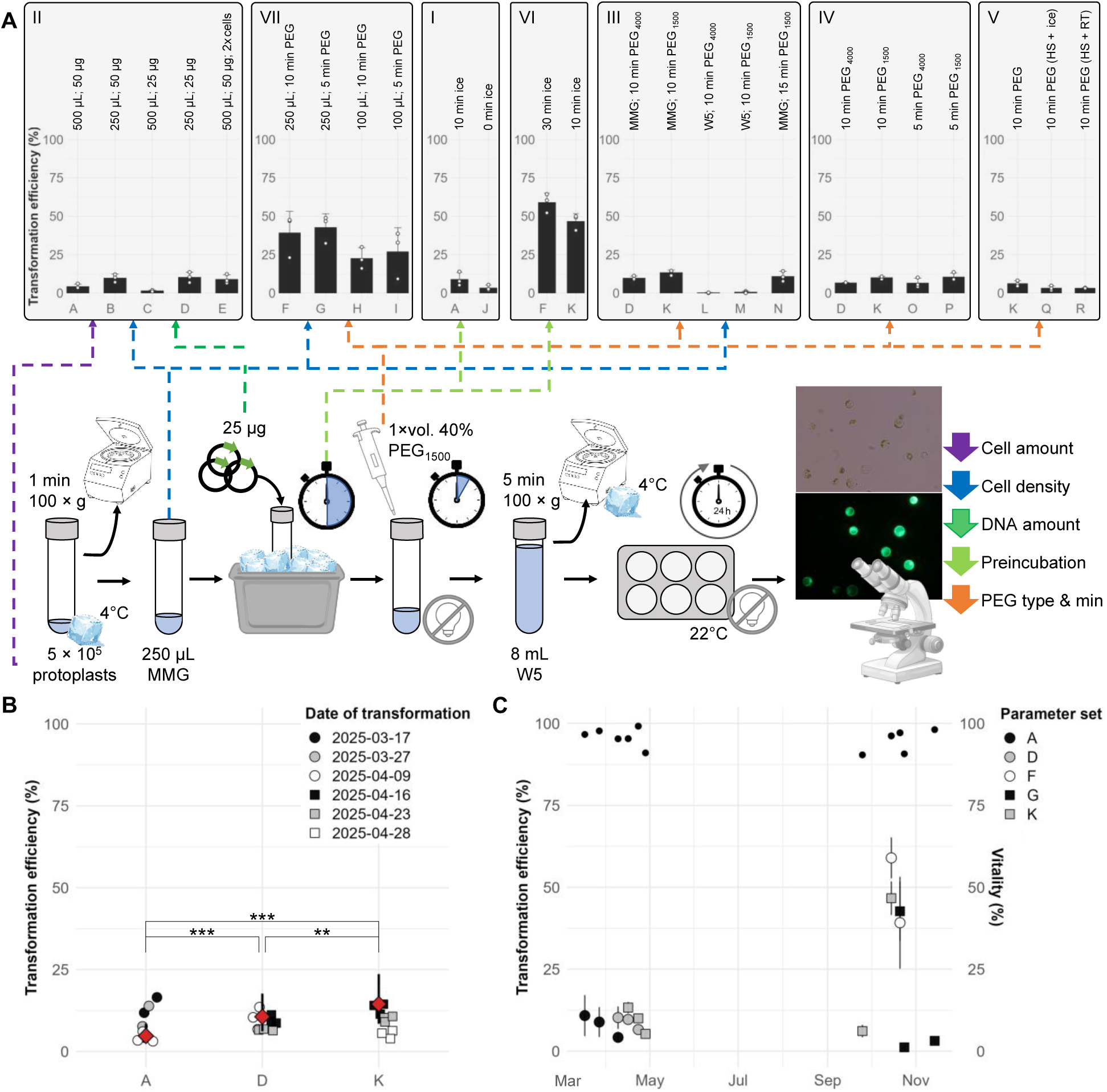
Effect of experimental setup and transformation date on PEG-mediated gene delivery in beech protoplasts. Protoplasts were transformed with the 35S::*mEGFP* reporter using parameter sets A–R (Table 2). Transformation efficiency (%) was defined as the proportion of GFP-positive cells relative to the total cell count (mean ± SD of three technical replicates). See Data S1 and Table S3 for numerical values (A) Overview of optimization experiments (I–VII; chronological order). Dot and bar plots illustrate raw transformation efficiencies and corresponding means ± SD for each parameter set. Parameter sets are grouped according to their position within the workflow and indicated by color-coded arrows. Key variables relevant to each experimental comparison are summarized above each bar, whereas shared parameters within a given comparison are not repeated. For complete experimental settings and parameter combinations, see Table 2. The optimized workflow (parameter set G) is shown schematically. All experiments used 5 × 10r protoplasts, except parameter set E (1 × 10ª; 2 × cells). Protoplasts were centrifuged at 100 × g for 1 min and resuspended in 500, 250, or 100 µL of W5 or MMG. Plasmid DNA (25 or 50 µg) was added and incubated on ice for 0, 10, or 30 min. Transformation was induced by adding one volume of 40% PEG (PEG_4000_ or PEG_1500_) and incubating for 5, 10, or 15 min in the dark. Alternatively, after PEG addition, heat shock (HS) was performed at 42°C for 1 min followed by 9 min on ice or at RT. Reactions were diluted to 8 mL with W5, centrifuged (5 min, 100 × g, 4°C), and incubated for 24 h at 22°C in the dark. Transformation efficiency was quantified via fluorescence microscopy. (B) Distribution and statistical comparison of transformation efficiencies for parameter sets A, D and K across biological replicates. Violin plots display raw efficiencies from March and April; points are color-coded by transformation date. Black dots represent Estimated Marginal Means from the GLMM; error bars show 95% confidence intervals. Asterisks indicate significant pairwise differences (*p & 0.05; **p & 0.01; ***p & 0.001). (C) Seasonal variation in transformation efficiency and protoplast vitality. Mean efficiencies (± SD) of parameter sets tested at least twice are shown in relation to the transformation date. Protoplast vitality (%) after isolation (Evans Blue or FDA staining) is plotted as an independent data layer (black dots; right-hand y-axis), enabling comparison between cell viability and transformation performance.

**Table 1:** Effect of varying PEG concentrations and incubation times on transformation efficiency. Transformation experiments were performed on two independent transformation days (No.) with 5 × 10r protoplasts in 250 (VI) or 100 µL (VII) MMG, and 25 µg plasmid DNA per reaction, using a PEG_1500_ solution. A pre-incubation on ice for 10 (VI) or 30 min (VII) was applied before PEG treatment. Transformation efficiency represents mean ± SD of three technical replicates.

| No. | Parameter set | PEG treatment | Transformation efficiency (mean ± SD) [%] |
| --- | --- | --- | --- |
| VI | K | 10 min 40% PEG | 46.69 ± 5.08 |
| VI | S | 20 min 20% PEG | 7.42 ± 0.70 |
| VII | H | 10 min 40% PEG | 22.54 ± 6.98 |
| VII | I | 5 min 40% PEG | 26.90 ± 15.59 |
| VII | T | 40 min 30% PEG | 12.80 ± 3.04 |
| VII | U | 40 min 20% PEG | 6.91 ± 4.33 |
| VII | V | 20 min 30% PEG | 13.11 ± 2.96 |

**Table 1:** WT-corrected modification rates and transformation-normalized editing efficiencies of *FsPDS*-targeting crRNAs based on deletion frequencies within ±10 bp of the predicted cut site determined using CLC-GWB. Modification rates were calculated from deletion-supporting reads relative to mean on-target coverage. WT background modification rates were subtracted before normalization to transformation efficiency.

| crRNA | crRNA 3_1 | crRNA 3_2 | crRNA 5_1 | crRNA 5_2 |
| --- | --- | --- | --- | --- |
| Mean coverage | 25193 | 24427 | 26133 | 29965 |
| Reads with deletion | 1427 | 365 | 218 | 249 |
| Modification rate (%) | 5.66 | 1.49 | 0.83 | 0.83 |
| WT-normalized (%) | 5.65 | 1.47 | 0.82 | 0.82 |
| Transformation efficiency (%) | 17.39 | 14.34 | 17.39 | 14.34 |
| Editing rate (%) | 32.69 | 10.27 | 4.75 | 5.71 |

**Table 2:** Parameter sets tested during the optimization of the PEG-mediated transformation protocol. . In all transformations, 5 × 10^5^ protoplasts were used, except for E, where 1 × 10^6^ protoplasts were used. For the heat shock (HS), cells were incubated for 1 min at 42°C and 9 min at room temperature (RT) or on ice. Each parameter set was used on at least one transformation day during the optimization (No.).

| Name | Transfection buffer | DNA [$\mu\text{g}$] | Ice [min] | PEG type | PEG [%] | PEG [min] | No. |
| --- | --- | --- | --- | --- | --- | --- | --- |
| A | 500 $\mu\text{L}$ MMG | 50 | 10 | 4000 | 40 | 10 | I, II |
| B | 250 $\mu\text{L}$ MMG | 50 | 10 | 4000 | 40 | 10 | II |
| C | 500 $\mu\text{L}$ MMG | 25 | 10 | 4000 | 40 | 10 | II |
| D | 250 $\mu\text{L}$ MMG | 25 | 10 | 4000 | 40 | 10 | II, III, IV |
| E | 500 $\mu\text{L}$ MMG | 50 | 10 | 4000 | 40 | 10 | II |
| F | 250 $\mu\text{L}$ MMG | 25 | 30 | 1500 | 40 | 10 | VI, VII |
| G | 250 $\mu\text{L}$ MMG | 25 | 30 | 1500 | 40 | 5 | VII |
| H | 100 $\mu\text{L}$ MMG | 25 | 30 | 1500 | 40 | 10 | VII |
| I | 100 $\mu\text{L}$ MMG | 25 | 30 | 1500 | 40 | 5 | VII |
| L | 250 $\mu\text{L}$ W5 | 25 | 10 | 4000 | 40 | 10 | III |
| K | 250 $\mu\text{L}$ MMG | 25 | 10 | 1500 | 40 | 10 | III, IV, V, VI |
| M | 250 $\mu\text{L}$ W5 | 25 | 10 | 1500 | 40 | 10 | III |
| N | 250 $\mu\text{L}$ MMG | 25 | 10 | 1500 | 40 | 15 | III |
| O | 250 $\mu\text{L}$ MMG | 25 | 10 | 4000 | 40 | 5 | IV |
| P | 250 $\mu\text{L}$ MMG | 25 | 10 | 1500 | 40 | 5 | IV |
| J | 500 $\mu\text{L}$ MMG | 50 | 0 | 4000 | 40 | 10 | I |
| Q | 250 $\mu\text{L}$ MMG | 25 | 10 | 1500 | 40 | 10 (HS + ice) | V |
| R | 250 $\mu\text{L}$ MMG | 25 | 10 | 1500 | 40 | 10 (HS + RT) | V |
| S | 250 $\mu\text{L}$ MMG | 25 | 10 | 1500 | 20 | 20 | VI |
| T | 250 $\mu\text{L}$ MMG | 25 | 30 | 1500 | 30 | 40 | VII |
| U | 250 $\mu\text{L}$ MMG | 25 | 30 | 1500 | 20 | 40 | VII |
| V | 250 $\mu\text{L}$ MMG | 25 | 30 | 1500 | 30 | 20 | VII |

Transformation efficiencies of the same parameter set showed considerable variability across independent experiments. In some cases, improved parameter sets yielded lower absolute efficiencies than suboptimal conditions from earlier time points (Figure 3A I, II). Initial analysis of parameter sets A, D, and K (n = 3 biological replicates) resulted in mean efficiencies of 7.98% ± 4.91%, 8.84% ± 2.49%, and 9.53% ± 3.67%, respectively. A one-way ANOVA revealed no significant differences in efficiency among the three groups (p = 0.847; Figure S4). However, in March and April, we observed a distinct pattern where efficiency for the same parameter set consistently decreased over time (Figure 3B, C). This temporal variation indicated a strong random effect of the transformation day on the outcome, necessitating a different statistical approach. To account for this variation, a GLMM was applied, including the transformation day as a random factor. A likelihood-ratio test revealed a significant overall effect of the parameter sets on transformation efficiency (χ² = 29.67, df = 2, p ≤ 0.001). Post-hoc analysis using the EMMs with Tukey adjustment confirmed clear differences: parameter set K (14.48%, 95% CI: 8.48–23.64%) showed significantly higher efficiency than D (10.62%, 95% CI: 6.17–17.68%), which in turn outperformed A (4.74%, 95% CI: 2.60–8.51%; Figure 3B). Model diagnostics indicated a generally acceptable fit (p > 0.05), although the uniformity test (p = 0.088) suggested slight deviations from the expected distribution, likely due to a lack of homogeneity for specific transformation days (23/04/25 and 28/04/25; Figure S5).

The first transformation in September yielded an efficiency comparable to the result obtained in April. The peak transformation efficiency was reached in October, with up to 59.00 ± 6.19%, followed by a sharp decline to 1.19 ± 0.87% (Figure 3C). The vitality of the protoplasts after isolation did not follow the same trend as the transformation efficiencies over the course of the year. Altogether, sequential optimization defined parameter set G as a suitable PEG-mediated transformation protocol for beech, despite a significant temporal effect on transformation efficiency.

### Identification of target gene *FsPDS*

Blasting the *Populus trichocarpa PDS* (*PtPDS*) gene sequence against the beech reference genome resulted in the identification of a putative *FsPDS* gene (Bhaga_5.g3100). A subsequent BLAST search using the putative *FsPDS* gene revealed the highest sequence similarity to *PDS* genes from *Fagaceae* family members, particularly to *Quercus robur PDS* (*QrPDS*), which served as a reference for subsequent structural and functional analyses (Table S4). The predicted *FsPDS* consists of 14 exons (Figure 4A; Figure S6a). An additional *in silico* predicted exon introducing a premature stop codon was excluded from the final gene model (Figure S6a). Exon 1 was added to the gene model by sequence homology to the *Fagaceae* species and was validated by the identified N-terminal chloroplast transit peptide (Figure S6c). The resulting gene structure includes the chloroplast translocation peptide and an amino oxidase domain architecture, which was found to be consistent with the highly homologous *Qr*PDS protein and the initial query sequence, *Pt*PDS (Figure S6b, c).

**Figure 4:**
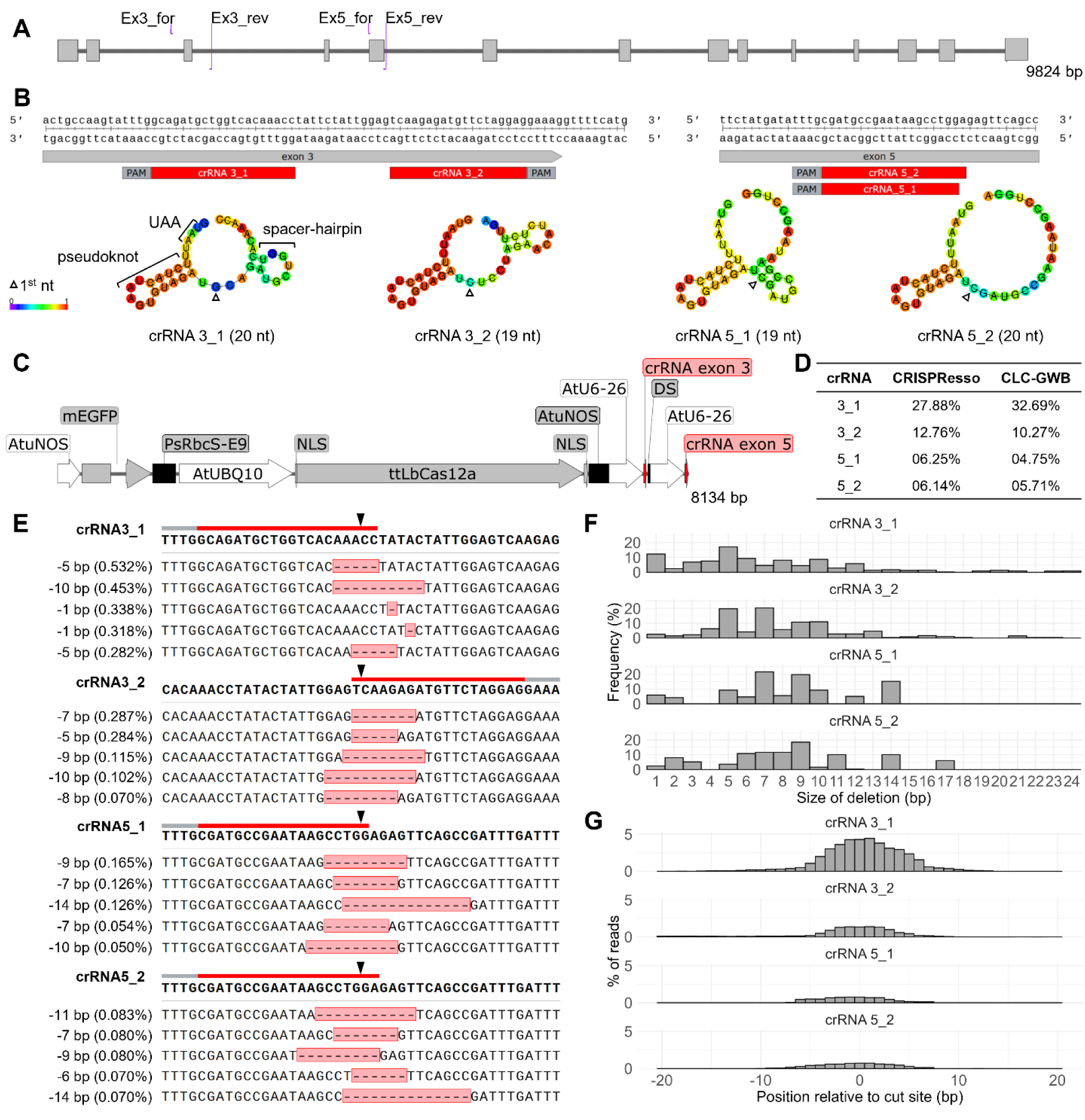
CRISPR/Cas12a-mediated editing of the *PHYTOENE DESATURASE* (*FsPDS*) gene in beech. (A) The *FsPDS* gene consists of 14 exons. Positions of the PCR primers used for target amplification are indicated (Ex3_for, Ex3_rev; Ex5_for, Ex5_rev). (B) Target regions within exon 3 and 5 are shown in magnified views, with the crRNA spacers (red) and the PAM (grey). *In silico* predicted secondary (centroid) structures of the crRNAs are shown with color-coded base pairing probabilities. The first target-determining nucleotide for each crRNA spacer is indicated. Structural features associated with crRNA efficiency are exemplarily highlighted for crRNA 3_1. The crRNAs ending on 1 are part of the FsPDS-KO1 vector, and crRNAs ending on 2 were encoded by the FsPDS-KO2 vector. (C) Partial schematic map of the vectors (FsPDS-KO1 and FsPDS-KO2) used for protoplast transformation. Each vector contains four expression cassettes consisting either of a coding sequence (grey) or a crRNA transcriptional unit (red), each flanked by a specific promoter (white) and terminator (black): (1) monomeric enhanced green fluorescent protein (mEGFP), (2) temperature-tolerant *Lb*Cas12a (ttLbCas12a) with two nuclear localization signals (NLS), (3+4) crRNA targeting exon 3 and exon 5 of *FsPDS* separated by a termination sequence (DS). (D) Comparison of transformation-normalized editing efficiencies for individual crRNAs calculated using CRISPResso 2.0 or CLC-GWB in consideration of transformation efficiencies. (E-G) CLC-GWB-based deletion analysis for each crRNA targeting *FsPDS*. (E) Top 5 deletion patterns. PAM (grey bar), spacer sequences (red bars), and cut sites (black triangles) are indicated above the WT reference sequence (first row). The frequency of each deletion pattern is shown relative to total read coverage and was not normalized to transformation efficiency. (F) Distribution of deletion sizes, shown as the frequency (%) of each deletion size within all deletions. (G) Distribution of deletions within ±20 bp of the cut site, expressed as a percentage of total reads.

### CRISPR/Cas-mediated editing of *FsPDS*

The modeled secondary structures of the gRNAs targeting *FsPDS* exons 3 and 5 predicted proper pseudoknot folding and absence of strong spacer internal base-pairing (Figure 4B). CCTop predicted medium editing efficiencies for both crRNA3_1 (0.66) and crRNA5_2 (0.62). To validate the functionality of the CRISPR/Cas12a system in beech, two distinct vectors, FsPDS-KO1 and FsPDS-KO2 targeting *FsPDS* with two gRNAs each (Figure 4C), were delivered into beech protoplasts using PEG-mediated transformation with mean transformation efficiencies of 12.92 ± 3.81% (FsPDS-KO1) and 9.79 ± 4.05% (FsPDS-KO2) (Table S5; Figure S7a, b). The presence of the *ttLbCas12a* gene was confirmed via PCR in all transformed samples.

Quantitative analyses of the amplicon sequencing yielded gene editing efficiencies in transformed cells ranging from 4.75% to 32.69%, depending on crRNA and quantification method (Table 3; Figure 4D). Table 3 additionally provides the corresponding overall modification rates across the total protoplast population. Top deletion patterns were unique for each crRNA used (Figure 4E). Deletion sizes ranged from 1 to 24 bp. Most frequent deletion sizes were 5 bp (crRNA3_1), 7 bp (crRNA3_2, crRNA5_1) and 9 bp (crRNA5_2). Across all four crRNAs, deletion events were concentrated in the range of 5–10 bp (Figure S8). Maximum deletion sizes were 24 bp (crRNA3_1), 21 bp (crRNA3_2), 14 bp (crRNA5_1) and 17 bp (crRNA5_2) (Figure 4F). In the WT, only a minor number of 1- and 2-bp deletions were found (Table S6). Subsequent deletion analysis revealed clear deletion peaks at the expected cleavage sites for all four targets (Figure 4G), confirming target-specific *FsPDS* editing. No insertions were found at the cut sites.

## Discussion

### Protoplast isolation and cultivation

We present a reproducible protocol for the isolation of beech protoplasts with high yield, which achieves excellent viability. This provides a robust foundation for functional studies in this woody species. The established system enables diverse transient cell-based applications, including subcellular localization, promoter activity assays, protein–protein interaction analyses, stress-response studies, and rapid screening of genome editing constructs prior to stable transformation. In addition, protoplasts provide a valuable platform for the functional characterization of candidate genes identified through genomic and transcriptomic approaches (Xu et al. 2022). However, protoplast-based assays remain limited to transient single-cell analyses, and the absence of a fully established regeneration system currently restricts the generation of stable edited plants from transformed beech protoplasts.

Seasonal and developmental factors strongly influenced isolation success. Highest yields were obtained from soft primary leaves shortly after emergence, during late winter to early summer and in autumn. Rapid leaf hardening outside these periods markedly reduced efficiency, even under constant growth conditions. This pattern is consistent with suggestions for seeding, seasonally programmed rhythms reported for beech seedling development, as well as seasonal variation in protoplast yield in other species, even when *in vitro* plants were used (Bonner and Leak, 2008; Keskitalo, 2001; Zahn et al., 2025). The persistence of these seasonal effects under controlled growth conditions further suggests regulation by endogenous circannual or epigenetically maintained developmental programs (García-Plazaola et al., 2017). Such physiological rhythms may not only affect protoplast isolation efficiency but also influence downstream transformation performance, consistent with the strong temporal variation in transformation efficiency observed in this study. Since each seedling batch consisted of a unique set of genotypes, genotype effects could not be separated from seasonal effects. Moreover, reduced germination success during summer may have resulted in a non-random subset of genotypes contributing to downstream phenotypic variation. The negative effect of leaf hardening on isolation is consistent with earlier reports for beech and other woody species (Ahuja, 1984; Han et al., 2023). Developmental changes in epicuticular wax composition during beech leaf maturation may accompany these seasonal effects and reflect broader physiological changes associated with leaf hardening (Gülz et al., 1992). Aligning isolation with these periods enabled yields of up to 1.5 × 108 protoplasts g⁻¹ FW with high viability. These values fall within the upper range reported for woody species and extend material availability in beech beyond the narrow spring flush typically used, while avoiding the labor-intensive and variable outcomes associated with forced bud break (Ahuja, 1984; Lang and Kohlenbach, 1988; Yang et al., 2024). Observed variability across replicates highlights the sensitivity of the procedure to leaf physiology and genotype, consistent with prior reports in plant protoplast systems (Ahuja, 1984; Yang et al., 2022). Manipulation of seed dormancy and physiological state, for example, through modified stratification regimes and storage gibberellic acid treatment, helps reduce season-dependent variability (Bonner and Leak 2008; Staszak et al. 2019). However, the extent to which these approaches stabilize seedling physiology in European beech remains unclear. Contamination control and antifungal sensitivity are critical. Mild surface sterilization combined with antimicrobial agents maintained low contamination and high viability. However, when added to the cultivation medium, natamycin caused rapid protoplast death, likely through non-specific interactions with membrane sterols (Hsueh et al., 2007; te Welscher et al., 2008), emphasizing the need for careful antifungal selection in wall-less plant cells.

The presence of chloroplast-free but viable protoplasts reflects inherent cellular heterogeneity, which is known in leaf-derived protoplast populations (Han et al., 2023) and should be considered in downstream experiments. Rapid cell wall regeneration and early proliferation indicate high physiological competence, supporting the improved initial regeneration compared with earlier reports in beech (Ahuja, 1984; Reed and Bargmann, 2021). However, cell proliferation ceased after microcallus formation, although no substantial cell death was observed. This suggests that the applied culture conditions supported early regeneration processes but remained inadequate for sustained callus proliferation and organogenesis, requiring further optimization of medium and culture conditions. In addition, the use of heterogeneous seed-derived material may further complicate regeneration, as regeneration competence in protoplast systems is often strongly genotype dependent and frequently restricted to only a subset of genotypes. Consequently, future optimization of beech protoplast regeneration may benefit from the establishment of defined regeneration-competent genotypes or clonal lines. This also eliminates potentially genotype-dependent effects on protoplast isolation and enables more precise evaluation of seasonal effects. Overcoming these regeneration bottlenecks will be essential for the future application of genome editing to functional studies of climate resilience in beech, including the validation of candidate genes associated with drought tolerance, stress adaptation, or pathogen resistance. In woody perennials, the ability to regenerate edited plants remains one of the major limiting factors for translating transient editing systems into stable functional genomics and breeding applications (Reed and Bargmann, 2021; Wang et al., 2025).

### Promoter-reporter validation

The functionality of six constitutive promoters (*CaMV*35S, *Atu*NOS, *Atu*OCS, *At*UBQ10, *Ha*UBI, *Pc*UBI) was successfully confirmed via transient expression using mEGFP or turboRFP reporters in beech protoplasts. The observed differences in the apparent transformation efficiency were likely attributed to two factors: promoter-dependent transcriptional activity and reporter-specific detection sensitivity. In particular, the turboRFP reporter is characterized by rapid maturation and higher intrinsic brightness compared to mEGFP (Merzlyak et al., 2007), which may facilitate detection of weakly expressing transformed cells. Therefore, the observed differences likely reflect both promoter-dependent expression levels and reporter-specific detectability rather than absolute promoter strength alone. Consistent with present data, the *UBIQUITIN* promoters generally drive higher gene expression compared to *CaMV*35S, whereas the *Atu*NOS and *Atu*OCS promoters are weaker (Horstmann et al., 2004; Kishi-Kaboshi et al., 2019; Wolabu et al., 2020). In addition, plasmid size has previously been reported to influence PEG-mediated transformation efficiency (Zetsche et al., 2015). In the present study, plasmid size ranged from 5,698 bp (*Atu*NOS::*mEGFP*) to 6,987 bp (*At*UBQ10::*mEGFP*), but did not appear to correlate with transformation efficiency. For example, 35S::*mEGFP* (6,937 bp) yielded higher transformation efficiencies than the substantially smaller *Atu*NOS::*mEGFP* (5,698 bp) and *Atu*OCS::*mEGFP* (5,872 bp) constructs, suggesting that promoter activity and reporter characteristics had a stronger influence on transformation outcomes than plasmid size within the tested range. In addition, variation in plasmid copy number between individual protoplasts may further influence detectable fluorescence levels. Consequently, rigorous quantitative comparisons of promoter strength will require future analyses using standardized dual-reporter systems with internal transformation controls and quantitative approaches such as qPCR-based transcript analyses, luciferase assays, or flow cytometry. Nevertheless, the confirmed functionality of these diverse promoters establishes the foundational molecular toolbox necessary for future transgenic studies in beech.

### PEG-mediated transformation and optimization

We successfully optimized the PEG-mediated transient transformation protocol for beech, achieving a peak efficiency of 59% (Figure 3) and establishing a reliable basis for transient expression assays.

Pre-incubation of DNA with protoplasts on ice for 30 min in Ca²⁺-containing MMG buffer was critical for maximizing efficiency at the reduced PEG incubation time (5 min), likely by promoting transient DNA–membrane interactions while protecting the plasmid from nucleases (Folling et al., 1998; Kinraide and Wang, 2010).

Transfection conditions were critical: MMG buffer (15 mM CaCl₂) outperformed W5 (125 mM CaCl₂), highlighting the importance of adequate cation concentration for DNA–membrane bridging and complex stabilization (Kinraide and Wang, 2010; Masani et al., 2014; Yoo et al., 2007). Doubling cell density from 1 × 10ª cells/mL to 2 × 10ª cells/mL markedly increased transformation efficiency, likely due to closer cell–DNA contact. Further increases in DNA (from 25 to 50 µg) or cell density (up to 1 × 10T cells/mL) did not improve efficiency, an effect also observed in woody plants, which may reflect saturation of uptake sites or crowding at high cell density (Masani et al., 2014; Zhao et al., 2025).

Beech protoplasts are highly sensitive to osmotic and chemical stress. PEG_1500_ consistently yielded higher transformation efficiencies than PEG_4000_, which may reflect lower cytotoxicity associated with lower molecular weight PEG (Hong et al., 2023). Although rarely used in plant protocols, it appears suitable for PEG-sensitive species (Bruegmann et al., 2019b). DNA uptake saturates within minutes, making the 5-min PEG incubation critical. Longer exposure likely reduces efficiency due to stress. In other species, optima can reach 30 min under the same PEG concentration (Bai et al., 2020), highlighting the high PEG sensitivity of beech. High PEG concentrations (40%) were necessary to achieve efficient DNA delivery (Table 1), whereas species with higher baseline efficiencies can suffice with lower, less cytotoxic concentrations (20%) (Yang et al., 2022). Heat shock, sometimes applied in other woody plants, proved detrimental in beech, suggesting that beech protoplasts are highly sensitive to additional stress during PEG-mediated transfection (Zhou et al., 2024).

Despite protocol optimization, transformation efficiency in beech remained consistently low (approx. 10 %) compared to model systems (e.g., *Populus* 80 %) or other non-model forest tree species (e.g., *Fraxinus excelsior* 67%, *Quercus ilex* 62%) (Lu et al., 2025; Pavese et al., 2024; Tan et al., 2013). High cell viability and effective cell wall removal confirmed that the low baseline was not due to technical failure. Instead, a major limiting factor was marked temporal variability across biological replicates, with efficiencies strongly correlated with the date of transformation. Temporal variation was accounted for in statistics, but residual differences may reflect external factors such as water stress (Yoo et al., 2007).

These fluctuations suggest an underlying endogenous, circannual rhythm, consistent with our observations in beech protoplast isolation and seedling responses to *in vitro* conditions (Zahn et al., 2025). This rhythm likely influences the protoplasts’ physiological response to transformation, affecting PEG stress tolerance and DNA uptake. Consistent with this, a 5-min PEG incubation was essential even under optimal conditions, as DNA uptake saturates rapidly and longer exposure reduces efficiency. Species-specific recalcitrance and heterogeneous genotypes may also contribute to the low baseline efficiency (Yang et al., 2022). It should also be noted that transformation efficiency was determined manually by fluorescence microscopy using only a subset of the total protoplast population (<0.19% of all cells per sample). Consequently, the reported efficiencies may be subject to observer-dependent variation and sampling bias. More precise quantification approaches, such as fluorescence-based flow cytometry or fluorescence-activated cell sorting (FACS), could further improve accuracy and throughput in future studies.

Although transformation efficiencies of approx. 50% are often considered desirable for downstream applications (Yoo et al., 2007), achieving a baseline efficiency of around 10% represents an important first milestone for beech, for which no protoplast transformation system has previously been established. While transformation efficiency showed pronounced temporal variability, the present work provides a reproducible and experimentally accessible framework for transient expression assays. Further optimization should focus on improving temporal stability. Overall, this protocol lays a solid foundation for functional studies and genome editing in beech and substantially expands the experimental toolbox for this species.

### CRISPR/Cas-mediated editing of *FsPDS*

The successful detection of deletions at all four target sites demonstrates the functionality of the tt*Lb*Cas12a system in beech protoplasts, a notable finding given the limited reports of Cas12a in woody plants (Das et al., 2025; Jia et al., 2022). Editing efficiencies varied between target sites, reaching up to 32.7 % for crRNA3_1, similar to stable poplar transformations with (tt)*Lb*Cas12a (An et al., 2020; Das et al., 2025). This is particularly noteworthy given that editing efficiencies in transient protoplast systems are typically much lower than in stable lines (Gao et al., 2015; Zhang et al., 2023).

Observed differences in crRNA efficiency align with literature on crRNA sequence- and secondary structure-dependent effects (Creutzburg et al., 2020; Kim et al., 2016). Our design prioritized proper pseudoknot formation, yet modeling revealed variations in spacer hairpins and base-pairing, likely influencing activity (Creutzburg et al., 2020). Minimal seed-region pairing (crRNA3_1) was tolerated, whereas stronger pairing (crRNA5_1) may reduce cleavage efficiency (Creutzburg et al., 2020; Zetsche et al., 2015). Other factors, such as UU motifs, spacer pairing position, hairpin loop size, and modeling uncertainties (e.g., exclusion of the poly-U tail), further complicate the prediction of crRNA efficiency (Creutzburg et al., 2020; Zhu and Liang, 2019). Interestingly, the two crRNAs targeting exon 5, which shared essentially the same target site and differed only in spacer length by 1 nt, showed comparable editing efficiencies. This may reflect locus-dependent effects on Cas12a activity and/or similarly favorable secondary structure configurations of the respective crRNAs (Bernabé-Orts et al., 2019). In addition, computational prediction tools such as CCTop were insufficient to reliably rank crRNAs. Together, these observations highlight the necessity of empirical validation, particularly in perennial species where stable transformation is time-consuming. Transient protoplast assays provide a rapid *in vivo* platform to assess crRNA efficacy and identify highly active guides before stable transformation (Naim et al., 2020; Panda et al., 2024; Sagarbarria et al., 2023).

Editing of *FsPDS* using tt*Lb*Cas12a resulted exclusively in deletions ranging from 1 to 24 bp, with distinct, crRNA-dependent mutational patterns, consistent with observations in poplar and pummelo (An et al., 2020; Das et al., 2025; Jia et al., 2022). Across the four crRNAs, deletions predominantly clustered between 5–10 bp, in line with Cas12a editing in other plants (Bernabé-Orts et al., 2019; Ma et al., 2024). Deletions of 1 bp, except for crRNA 5_1, might be considered sequencing artifacts. Editing efficiencies estimated by CRISPResso and manual analyses were highly similar, preserving the relative ranking of crRNAs, though manual analysis captured complex or low-frequency deletion patterns more effectively. The efficient PCR-based approach did not consider editing events spanning both Cas12a target site, because exons 3 and 5 were amplified and analyzed separately. Generally, multiplexed Cas12a approaches can cause deletions between paired target sites which would amend the detected editing patterns (Wang et al., 2017).

Although *PDS* is widely used as a visual marker gene due to the albino phenotypes observed in regenerated tissues and plants, no obvious bleaching phenotype was detected in the present transient protoplast assays. This is likely attributable to several factors. First, despite relatively high editing efficiencies among transformed cells, the overall proportion of edited cells within the total protoplast population remained low due to the limited transformation efficiency. Second, the transient assay duration (24–48 h) was likely insufficient for substantial carotenoid depletion and subsequent chlorophyll degradation to become visibly apparent, particularly in non-dividing cells (Hirschberg, 2001). Finally, the presence of chloroplast-free yet viable protoplasts further reduced the proportion of cells capable of developing detectable pigment-related phenotypes. Consistent with previous studies, visible *PDS*-associated albino phenotypes are typically observed only after sustained cell proliferation and regeneration of edited tissues or plants (Pavese et al., 2021). Because the present study focused on establishing a proof-of-concept transient editing platform, crRNAs were selected manually based on target suitability and predicted secondary structure rather than comprehensive genome-wide off-target evaluation. Potential off-target activity was therefore not experimentally assessed and cannot be excluded, particularly given the heterozygosity and still incompletely validated annotation of the beech genome (Mishra et al., 2021; Stemmer et al., 2015).

Overall, these results demonstrate that tt*Lb*Cas12a induces a characteristic, target-site- and crRNA-dependent deletion spectrum in beech, closely matching outcomes in other woody and herbaceous species. Our findings confirm the suitability of the presented system for rapid validation of multiplex genome editing constructs and emphasize the importance of empirical crRNA validation in perennial plants. However, the application of this platform to stable functional studies in beech will ultimately depend on further advances in regeneration systems and regeneration-competent lines.

## Conclusions

The optimization of protoplast isolation and transient transformation for beech provides a robust, high-yield system, achieving up to 1.5 × 108 protoplasts g⁻¹ FW and a peak transient efficiency of 59%. This was accomplished by addressing key biological constraints, including the need to align preparations with the endogenous circannual rhythm and by developing protocols that accommodate the pronounced PEG sensitivity of beech protoplasts. We further established a foundational, MoClo-compatible molecular toolbox using functional promoters and demonstrated the first successful application of tt*Lb*Cas12a-mediated genome editing in beech, reaching editing efficiencies of up to 32.69%. However, the strong temporal variability in transformation efficiency and the additional genotype-dependent variation introduced by seed-derived material indicate that further optimization is required to improve reproducibility. Ongoing advances in beech *in vitro* culture may facilitate the future establishment of standardized regeneration-competent lines (Thiesen et al., 2025; Zahn et al., 2025), which would further strengthen the utility of the presented protoplast platform as a rapid *in vivo* validation system for targeted genome editing and future functional genomics studies in woody species.

## Experimental procedures

### Plant material

Beech nuts were collected in autumn 2024 in the arboretum of the Thünen Institute, Grosshansdorf, Germany. The seeds were stored cool and dark until germination. Germination was initiated in darkness at 4°C in moist clay granulate until the radicles emerged (January to May and September to November 2025). Seedling development was carried out at 22°C under a 16-hour photoperiod (70 µmol m⁻² s⁻¹ photosynthetic photon flux density) in clay granulate in a climate chamber. The expansion of the primary leaves took approx. three weeks.

### Protoplast isolation

Protoplast isolation was performed using a protocol adapted from Bruegmann et al. (2019b). Unless stated otherwise, all experiments were performed using the finalized protocol. Primary leaves were harvested 24–48 h after unfolding (Figure 1A). Leaf samples (0.51–0.81 g) were washed in 1% Mucasol (alkaline detergent; Schülke & Mayr, Norderstedt, Germany) and surface-sterilized with 0.12% NaClO containing 0.1% Tween-20 for 10 min, followed by rinsing with sterile water. Cleaned leaves were cut into approx. 3 mm wide strips and washed 3 times for 30 min in basal solution (10 mM CaCl_2_, 0.2 mM KH_2_PO_4_, 1 mM KNO_3_, 1 mM MgSO_4_·7H_2_O, 5 mM MES, 1% PVP-10; adjusted to 600 mOsmol/kg with mannitol, pH 5.6).

The enzyme solution (2% (w/v) cellulase, 1% (w/v) macerozyme, 10 mM CaCl_2_, 1 mM KH_2_PO_4_, 1 mM MgSO_4_, 20 mM MES, 2% (w/v) PVP-40, and 0.25% (w/v) BSA; adjusted to 600 mOsmol/kg with mannitol, pH 5.6) was heated to 40°C for 10 min, and supplemented with antibiotics (500 mg/L cefotaxime, 50 mg/L gentamicin, 50 mg/L streptomycin) and fungicide (10 mg/L natamycin; HPC Standards, Borsdorf, Germany). All enzymes and antibiotics were supplied by Duchefa Biochemicals (Haarlem, The Netherlands).

25 mL of enzyme solution was added to the leaf strips before vacuum infiltration 3 times for 10 min at 7 mbar. Enzymatic digestion was carried out at 22°C in the dark for 16 h (4 rpm on an orbital shaker). The resulting suspension was filtered through a 30 µM nylon mesh (Lang and Kohlenbach, 1988). The remaining tissue was washed 3 times with 10 mL W5 solution (125 mM CaCl₂, 5 mM KCl, 5 mM glucose, 155 mM NaCl; pH 5.6) with repeated filtering. Protoplasts were collected by centrifugation at 50 × g for 6 min, and the pooled protoplasts were resuspended in W5 solution. Protoplast yield was quantified using a hemocytometer. In preliminary experiments, protoplast viability was assessed using Evans Blue (400 mg/L in 0.5 M mannitol), mixed 1:1 (v/v) with the protoplast suspension, whereas fluorescein diacetate (FDA) staining was used for all subsequent experiments (Bruegmann et al., 2019b).

### Protoplast culture

Protoplasts were cultured at a 2 × 10r cells/mL density in cultivation medium (Murashige and Skoog medium including vitamins, 0.01 µM thidiazuron (TDZ), 14 µM 2,4-dichlorophenoxyacetic acid (2,4-D); adjusted to 600 mOsmol/kg with glucose/mannitol (2:1), pH 5.6; modified from Nietsch et al. (2017)) in 6-well plates at 22°C in the dark to assess sterility of the isolation procedure and protoplast development, unless all protoplasts were used for transformation. All the medium ingredients were supplied by Duchefa Biochemicals. To evaluate the effect of antimicrobial agents during protoplast culture in initial experiments, the antibiotics only and the antibiotics in combination with the fungicide in the concentrations mentioned above were added to the cultivation medium. The medium was exchanged every 6 weeks. To monitor cell wall regeneration, 5 × 10^5^ protoplasts were suspended in 1 mL of cultivation medium and cultivated in a 10 mL round-bottom centrifuge tube at 22°C in the dark until cell wall staining.

### Cell wall staining

Staining of protoplasts, either to assess the presence of residual cell wall fragments after isolation or to visualize cell wall regeneration over time in cultivation medium, was performed using Calcofluor White. For staining, Calcofluor White (1 g/L; Sigma-Aldrich, St. Louis, MO, USA) was added to protoplast suspensions in W5 solution (after isolation) or in cultivation medium (after cultivation) at a final dilution of 1:10 (v/v) (Neubauer et al., 2022). After 2 min incubation at RT in the dark, the protoplasts were centrifuged at 60 × g for 3 min. The supernatant was discarded. The protoplasts were washed with W5 solution and finally resuspended in 50 µL W5. 20 µL of protoplasts were used for fluorescence microscopy.

### Cloning of promoter–reporter constructs

Expression vectors for promoter and reporter gene analysis in beech were assembled using the Modular Cloning (MoClo) system (Marillonnet and Grützner, 2020). Modules from the MoClo Plant Parts Kit were complemented with additional elements (Table S7). To ensure compatibility with MoClo, internal *Bsa*I and *Bpi*I recognition sites within the additional elements were removed by PCR-based site-directed mutagenesis (Table S8). All primers for cloning are listed in the supplements (Table S9).

Level-1 (L1) expression vectors were generated, including constructs expressing the monomeric Enhanced Green Fluorescent Protein (mEGFP) under the control of the CaMV35S promoter with the TMV-Ω 5’UTR (35S::*mEGFP*), the *Arabidopsis thaliana UBIQUITIN10* promoter including 5’UTR and first intron (*At*UBQ10::*mEGFP*), the *Helianthus annuus POLYUBIQUITIN Ha*UBI promoter (*Ha*UBI::*mEGFP*), the *Agrobacterium tumefaciens NOPALINE SYNTHASE* promoter (*Atu*NOS::*mEGFP*) and the *A. tumefaciens OCTOPINE SYNTHASE* promoter (*Atu*OCS::*mEGFP*). Transcription was stopped by *A. thaliana* HEAT SHOCK PROTEIN 18.2 (*At*HSP18.2) terminator. Additionally, a turboRFP construct with the *Petroselinum crispum UBIQUITIN* promoter and *Pisum sativum* pea3A terminator was generated (*Pc*UBI::*turboRFP*). All vector sequences (.gb files) are available in Data S2.

### Validation of genome editing target

Using BLAST analysis based on the *PDS* gene sequence from *PtPDS* (Potri.014G148700), the target gene was identified in beech using the reference genome from (Mishra et al., 2021). The target regions of *FsPDS* were additionally verified in northern German beech individuals by PCR amplification and subsequent Sanger sequencing (Table S9). The exon-intron structure was predicted *in silico* using FGENESH (Softberry Inc., http://www.softberry.com), employing *P. trichocarpa* as a reference. Predicted exons, the annotated *PtPDS* gene, and mRNA sequences of tree species from the *Fagaceae* family were aligned to the *FsPDS* sequence to support exon identification. Species were selected based on the highest homology using the NCBI BLAST search. Sequence alignments were made using SnapGene software (Dotmatics, Boston, MA, USA). Protein domains were annotated for the predicted FsPDS protein in SMART (Letunic et al., 2021), and signal peptides were screened using TargetP-2.0 (Almagro Armenteros et al., 2019). The resulting annotations were compared to protein sequences derived from *PtPDS* and *QrPDS* (belonging to the *Fagaceae* family) to support the structural and functional analysis.

### gRNA and editing vector design

CRISPR/Cas12a target sites were identified by scanning exon regions of the *FsPDS* gene for the canonical TTTV PAM. Spacer sequences (19–24 nt) complementary to the target DNA were selected. A non-canonical AsCas12a-derived direct repeat was used (Zetsche et al., 2015), which is compatible with *Lb*Cas12a (Nguyen et al., 2024). A 5’ G was added to support U6 promoter–driven transcription (Shan et al., 2013), and transcription termination was achieved using a 7×T tail (Verosloff et al., 2021). Secondary structures were predicted using RNAfold (Lorenz et al., 2011) with RNA parameters from (Andronescu et al., 2007) at 22°C. Candidate crRNAs were prioritized based on stable pseudoknot formation in the direct repeat, unpaired UAA motifs, minimal spacer self-complementarity, GC content between 30–70%, and absence of homopolymeric triplets (Creutzburg et al., 2020; Zhu and Liang, 2019). Two crRNAs each targeting Exon 3 and Exon 5 were selected (5’ – 3’): crRNA3_1 (gcagaugcuggucacaaacc), crRNA3_2 (ucaagagauguucuaggag), crRNA5_1 (cgaugccgaauaagccugg), and crRNA5_2 (cgaugccgaauaagccugga). Predicted on-target activities were assessed *in silico* using CCTOP (Stemmer et al., 2015). While the promoter constructs were assembled using the MoClo system, the final editing vectors FsPDS-KO1 and FsPDS-KO2 were commercially synthesized by DNA Cloning Service (Hamburg, Germany) due to their large size and repetitive sequence composition. Each vector contained an AtuNOS::mEGFP cassette, an AtUBQ10::ttLbCas12a cassette, and two AtU6-26::crRNA expression units targeting *FsPDS*. Fully annotated vector sequences (.gb files) are provided in Data S2. The weak NOS promoter was used to drive *mEGFP* expression, allowing prolonged incubation without detrimental cytotoxicity, while AtU6-26 and AtUBQ10 promoters were selected to ensure robust crRNA and *ttLbCas12a* expression, respectively, based on their reported efficacy in plant systems (Bruegmann et al., 2019a; Castel et al., 2019; Horstmann et al., 2004; Li et al., 2007; Lu et al., 2025; Walawage et al., 2019; Wolabu et al., 2020).

### PEG-mediated transformation

Protoplast transformation was performed using a PEG-mediated protocol adapted from (Bruegmann et al., 2019b). Freshly isolated protoplasts stored in W5 solution at 4°C were used. Key transformation parameters were optimized using a 35S::mEGFP reporter construct (Table 2). For the final protocol (parameter set G), 5 × 10r protoplasts were resuspended in 250 µL MMG solution (4 mM MES, 0.4 M mannitol, 15 mM CaCl₂, adjusted to 600 mOsmol/kg with glucose, pH 5.6) and incubated with 25 µL plasmid DNA (1 µg µL⁻¹; PureYield Plasmid Miniprep System; Promega, Madison, WI, USA) on ice for 30 min. PEG_1500_ (40%) was added at a 1:1 ratio, followed by incubation for 5 min in the dark. The reaction was terminated by stepwise addition of 8 mL W5 solution and centrifugation at 100 × *g* for 5 min at 4°C. Protoplasts were resuspended in cultivation medium at 2 × 10r mL⁻¹ and incubated in 6-well plates at 22°C in the dark for 24 or 48 h (CRISPR/Cas vectors only).

Reporter constructs were transformed using parameter sets A and G in two independent experiments; CRISPR/Cas vectors used parameter set K (50 µg DNA). Three technical replicates were performed per parameter set or vector. Transformation efficiency for all vectors was quantified by fluorescence microscopy as the percentage of mEGFP- or turboRFP-positive cells. For each replica, 71 to 944 cells were counted (Data S1). Values are reported as mean ± SD. For microscopy, protoplasts were released from plates by incubation with enzyme solution for 2 h at room temperature, pooled, centrifuged (100 × g, 5 min, 4°C), and resuspended for observation.

### Microscopy

Protoplast counting, viability assessment by Evans Blue, and morphological checks were performed using an inverted light microscope. Fluorescence microscopy of FDA- and Calcofluor White-stained protoplasts, as well as of transformed cells, was conducted on an Axioscope 3 fluorescence microscope (Carl Zeiss Microscopy, Jena, Germany). Filter sets were as follows: mEGFP and FDA 469/38 nm (Ex) / 520–550 nm (Em); turboRFP 555/30 nm (Ex) / 600–650 nm (Em); Calcofluor White 385/30 nm (Ex) / 445/50 nm (Em). ISO settings for transformed protoplasts were calibrated against PEG-treated wildtype (WT) protoplasts to confirm absence of autofluorescence. Image analysis and cell counting were performed using Fiji (Schindelin et al., 2012). Spatial calibration metadata were not recorded during image acquisition. All microscopy images within each experiment were acquired using identical magnification settings.

### Detection of genome editing events

Genomic DNA was extracted from protoplasts transformed with CRISPR/Cas vectors FsPDS-KO1 or FsPDS-KO2 (Bruegmann et al., 2022). The presence of *ttLbCas12a* was verified by standard PCR. Target regions within *FsPDS* Exons 3 (Ex3; 415 bp) and 5 (Ex5; 175 bp) were amplified by PCR with Q5 High-Fidelity DNA Polymerase (New England Biolabs, Ipswich, MA, USA). Primer sequences are listed in Table S9. The amplified target regions of the transformations with the highest transformation efficiency as well as the WT control, were purified with the QIAquick PCR Purification Kit (QIAGEN, Hilden, Germany) and individually submitted to GENEWIZ (Leipzig, Germany) for paired-end Illumina sequencing (2 × 250 bp). The fastq files from the amplicon sequencing are available in the European Nucleotide Archive (ENA; Accession No. PRJEB107651).

The raw FastQ files were used for downstream indel analysis with the cutting side set after the 18^th^ nucleotide of the spacer sequence. For CRISPResso 2.0, the quantification window was set at 10 bp, and substitutions were ignored (Clement et al., 2019). For further analysis of indels due to insufficient results from CRISPresso, Illumina amplicon sequencing reads from six datasets (FsPDS-KO1_Ex3/Ex5, FsPDS-KO2_Ex3/Ex5 and WT_Ex3/Ex5) were quality-trimmed and mapped to the edited exon 3 or exon 5 reference sequences (Table S10) using CLC Genomics Workbench (CLC GWB; QIAGEN).

Variants, including rare variants, were detected, and indels overlapping the target region (cut site ± 10 bp) were selected for further analysis. The modification rate (%) was calculated as the number of deletion-supporting reads divided by the mean coverage at the target site. Position-specific deletion frequencies were determined by summing all deletions spanning each nucleotide position. Reads with representative deletion patterns were selected based on BLAST analysis of sequences flanking the cut site (± 20 bp) versus trimmed reads. Detailed trimming, mapping, variant calling, and BLAST parameters are provided in Method S1.

Editing rates were calculated by subtracting background modification rates detected in WT controls from the observed modification rates and subsequently normalizing the values to transformation efficiency according to the following equation:

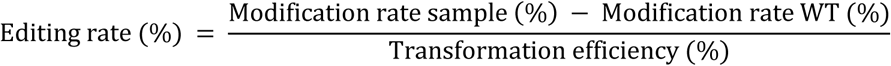

Since only a fraction of the total protoplast population was successfully transformed, normalization to transformation efficiency was used to estimate editing activity within the transformed cell fraction.

### Statistics and data visualization

Statistical analyses and data visualizations were performed in RStudio v2024.12.0.467. Significance was defined as p ≤ 0.05. Parameter optimization results (n = 3 technical replicates) are reported as mean ± SD. Transformation efficiency data (n = 3 biological replicates) were analyzed using one-way ANOVA and generalized linear mixed models (GLMMs) to account for nested structure and binomial count data. For GLMM analyses, temporally matched biological replicates were selected to minimize confounding effects caused by strong seasonal variation in transformation efficiency, while transformation date was included as a random factor. Model assumptions and diagnostics were assessed, and post hoc comparisons were performed using estimated marginal means (Tukey method). Details on statistical analyses are described in Method S2 and Data S3. For graphics with low percentage values, y-axis ranges were adjusted to improve visual resolution.

## Supporting information

Data S1 - Raw data transformation optimization

Data S2 - Vector and target sequences

Data S3 - Statistical data

## Authors’ Contributions

Study conception and design – V.Z., M.F., T.B.; methodology implementation – V.Z., A.-J.S., T.B.; experiment execution – V.Z., A.-J.S.; data collection – V.Z.; data analysis/interpretation – V.Z., B.K., T.B.; manuscript writing/revision – V.Z., B.K., M.F., T.B.; funding acquisition – M.F., T.B.

## Acknowledgments

We like to thank Susanne Jelkmann for her excellent laboratory assistance and Ina Martini for fluorescence assays. The work was supported by the German Federal Ministry for Agriculture, Food, and Regional Identity (BMLEH) via the Fachagentur Nachwachsende Rohstoffe within the project TreeEdit (grant 2219NR359). Open Access funding enabled and organized by Project DEAL. This article is based upon work from COST Action CA21157 “European Network for Innovative Woody Plant Cloning”, supported by COST (European Cooperation in Science and Technology), www.cost.eu.

## Data Availability

Plasmids generated and used in this study are available from the corresponding author upon reasonable request. Amplicon sequencing data have been deposited in ENA under accession number PRJEB107651.

## Supplementary Data

Method S1: Detection of genome editing events

Method S2: Statistics and data visualization

Figure S1: Beech protoplast viability after 5 (a-c) and 9 days (d-f) in different media

Figure S2: Negative control of protoplast transfection for promoter-reporter validation

Figure S3: Protoplast morphology and mEGFP expression of F. sylvatica protoplasts 24 h post-transformation using PEG1500

Figure S4: QQ-plot of ANOVA residuals for protoplast transformation efficiencies (parameters A, D, and K)

Figure S5: Diagnostic plots of GLMM for comparison of transformation parameters over time

Figure S6: Annotation of F. sylvatica PHYTOENE DESATURASE (FsPDS)

Figure S7: F. sylvatica protoplasts after transformation with the CRISPR/Cas12a vectors

Figure S8: Cumulative distribution of deletion sizes induced by ttLbCas12a across four crRNAs targeting FsPDS in F. sylvatica protoplasts

Table S1: Protoplast isolation from *F. sylvatica* seedling leaves

Table S2: Transformation efficiencies of different promoter reporter constructs (vector) used for *F. sylvatica* protoplast transformation

Table S3: Transformation efficiencies of different parameter sets used for *F. sylvatica* protoplast transformation

Table S4: Highest ranked results of BLAST search of *FsPDS* gene against the NCBI database

Table S5: Transformation efficiency of beech protoplasts using CRISPR/tt*Lb*Cas12a vectors

Table S6: Deletions detected within ±10 bp of the cut site for *FsPDS*-targeting crRNAs in the wildtype control

Table S7: Level-0 (L0) modules used for cloning of L1 promoter-reporter constructs.

Table S8: Modifications of Level-0 modules for MoClo compatibility

Table S9: Primers used in this study

Table S10: Edited reference sequences of *FsPDS* exons 3 and 5 used for alignment of amplicon sequencing data

Data S1: Raw data transformation optimization

Data S2: Vector and target sequences

Data S3: Statistical data

