## Supplemental Tables for "First report of genetic transformation and CRISPR/Cas12a-mediated gene editing of European beech (*Fagus sylvatica* L.) employing a transient protoplast system"

**Table S1: Protoplast isolation from *F. sylvatica* seedling leaves.** Yield is expressed relative to the amount of leaf material used—per fresh weight ($g^{-1}$ FW). Protoplast viability was assessed by Evans Blue (EB) or fluorescein diacetate (FDA) staining. The viability rate was calculated as the ratio of blue-colored or green-fluorescent cells to the total number of cells counted (n ≥ 240), respectively. The contamination rate was calculated as the ratio of contaminated wells to all wells (n = 6 or 12). When no contamination rate was assessed (X), no protoplasts remained for culture after the transformation experiments. Values highlighted in gray were not included in the calculation of average protoplast yield or viability, as a precursor protocol or an alternative staining method was used.

| Date of isolation | g FW | Protoplasts $g^{-1}$ FW | Staining method | Vitality rate (%) | Contamination rate (%) |
|---|---|---|---|---|---|
| 28/01/2025 | 0.81 | $3.12 \times 10^7$ | EB | 94.17 | 0.00 |
| 17/03/2025 | 0.68 | $3.57 \times 10^7$ | EB | 96.66 | 0.00 |
| 25/03/2025 | 0.55 | $5.13 \times 10^7$ | EB | 90.04 | 0.00 |
| 27/03/2025 | 0.61 | $10.98 \times 10^7$ | EB | 97.74 | 0.00 |
| 09/04/2025 | 0.81 | $1.77 \times 10^7$ | FDA | 95.32 | 0.00 |
| 16/04/2025 | 0.64 | $14.63 \times 10^7$ | FDA | 95.37 | 0.00 |
| 23/04/2025 | 0.53 | $3.90 \times 10^7$ | FDA | 99.19 | 0.00 |
| 28/04/2025 | 0.60 | $1.33 \times 10^7$ | FDA | 91.02 | X |
| 06/05/2025 | 0.52 | $4.16 \times 10^7$ | FDA | 88.76 | 16.67 |
| 25/09/2025 | 0.52 | $6.26 \times 10^7$ | FDA | 90.42 | 0.00 |
| 15/10/2025 | 0.51 | $3.98 \times 10^7$ | FDA | 96.22 | 0.00 |
| 21/10/2025 | 0.54 | $5.56 \times 10^7$ | FDA | 97.16 | 25.00 |
| 24/10/2025 | 0.53 | $3.78 \times 10^7$ | FDA | 90.73 | X |
| 14/11/2025 | 0.58 | $4.29 \times 10^7$ | FDA | 98.12 | 0.00 |

**Table S2: Transformation efficiencies of different promoter reporter constructs (vector) used for *F. sylvatica* protoplast transformation.** Efficiency was calculated by the mean of transformation efficiency (% GFP-positive cells relative to total cells) of the three technical replicates. Parameter sets are in accordance with Table 1.

| Date of transformation | Parameter set | Vector | Transformation efficiency (mean ± SD) [%] |
|---|---|---|---|
| 17/03/2025 | A | 35S::*mEGFP* | 10.86 ± 6.21 |
| 17/03/2025 | A | *Atu*NOS::*mEGFP* | 9.19 ± 2.37 |
| 17/03/2025 | A | *Ha*UBI::*mEGFP* | 15.03 ± 1.28 |
| 17/03/2025 | A | *Pc*UBI::*mEGFP* | 21.77 ± 2.02 |
| 24/10/2025 | G | 35S::*mEGFP* | 1.19 ± 0.87 |
| 24/10/2025 | G | *Atu*OCS::*mEGFP* | 0.45 ± 0.64 |
| 24/10/2025 | G | *At*UBQ10::*mEGFP* | 3.05 ± 1.24 |

**Table S3: Transformation efficiencies of different parameter sets used for *F. sylvatica* protoplast transformation.** Efficiency was calculated by the mean of transformation efficiency (% GFP-positive cells (cells_GFP) relative to total cells (cells_total)) of the three technical replicates. No. and parameter sets are in accordance with Table 1.

| Date of transformation | No. | Parameter set | cells_total | cells_GFP | Transformation efficiency (mean ± SD) [%] |
|---|---|---|---|---|---|
| 17/03/2025 | - | A | 378 | 45 | 10.86 ± 6.21 |
| 27/03/2025 | I | A | 878 | 84 | 8.89 ± 4.51 |
| 27/03/2025 | I | P | 2029 | 59 | 3.28 ± 1.83 |
| 09/04/2025 | II | A | 814 | 34 | 4.21 ± 1.63 |
| 09/04/2025 | II | B | 640 | 63 | 9.73 ± 2.54 |
| 09/04/2025 | II | C | 1204 | 16 | 1.45 ± 0.43 |
| 09/04/2025 | II | D | 753 | 81 | 10.25 ± 3.38 |
| 09/04/2025 | II | E | 493 | 44 | 8.91 ± 3.01 |
| 16/04/2025 | III | D | 1184 | 119 | 9.68 ± 1.33 |
| 16/04/2025 | III | J | 1360 | 3 | 0.18 ± 0.16 |
| 16/04/2025 | III | K | 1086 | 145 | 13.30 ± 1.65 |
| 16/04/2025 | III | L | 1148 | 6 | 0.56 ± 0.61 |
| 16/04/2025 | III | M | 908 | 105 | 10.79 ± 3.31 |
| 23/04/2025 | IV | D | 1072 | 71 | 6.60 ± 0.23 |
| 23/04/2025 | IV | K | 1167 | 119 | 10.02 ± 0.88 |
| 23/04/2025 | IV | N | 1253 | 82 | 6.49 ± 3.15 |
| 23/04/2025 | IV | O | 1022 | 104 | 10.45 ± 2.67 |
| 28/04/2025 | - | K | 1169 | 60 | 5.28 ± 1.00 |
| 25/09/2025 | V | K | 805 | 49 | 6.15 ± 1.84 |
| 25/09/2025 | V | Q | 806 | 25 | 3.18 ± 1.55 |
| 25/09/2025 | V | R | 684 | 21 | 3.14 ± 0.27 |
| 15/10/2025 | VI | K | 884 | 418 | 46.69 ± 5.08 |
| 15/10/2025 | VI | F | 772 | 454 | 59.00 ± 6.19 |
| 15/10/2025 | VI | S | 907 | 67 | 7.42 ± 0.70 |
| 21/10/2025 | VII | F | 892 | 350 | 39.20 ± 13.98 |
| 21/10/2025 | VII | G | 876 | 366 | 42.70 ± 9.01 |
| 21/10/2025 | VII | H | 1225 | 282 | 22.54 ± 6.98 |
| 21/10/2025 | VII | I | 1019 | 261 | 26.90 ± 15.59 |
| 21/10/2025 | VII | T | 1126 | 139 | 12.80 ± 3.04 |
| 21/10/2025 | VII | U | 1269 | 97 | 6.91 ± 4.33 |
| 21/10/2025 | VII | V | 1130 | 143 | 13.11 ± 2.96 |
| 24/10/2025 | - | G | 907 | 13 | 1.19 ± 0.87 |
| 14/11/2025 | - | G | 1046 | 88 | 3.16 ± 0.49 |

**Table S4: Highest ranked results of BLAST search of *FsPDS* gene against the NCBI database.** The *Populus* results are included for comparison.

| Species | Accession | Max score | Total score | Query cover (%) | E-value | Identity (%) | Alignment length |
|---|---|---|---|---|---|---|---|
| *Quercus suber* | XM_065777133.1 | 355 | 1984 | 13 | 8e-92 | 96.3 | 2426 |
| *Quercus robur* | XM_050436637.1 | 355 | 2523 | 17 | 8e-92 | 96.3 | 2330 |
| *Castanea sativa* | XM_075797933.1 | 350 | 2528 | 17 | 4e-90 | 95.8 | 2365 |
| *Quercus lobata* | XM_031066959.1 | 346 | 2368 | 16 | 5e-89 | 95.4 | 2384 |
| *Populus trichocarpa* | XM_002321068.4 | 235 | 686 | 6 | 5e-59 | 92.2 | 2195 |

**Table S5: Transformation efficiency of beech protoplasts using CRISPR/tt*Lb*Cas12a vectors.** Efficiency [%] was quantified as the percentage of green fluorescent cells relative to the total cell count. Mean efficiency is presented for each vector based on three independent technical replicates (Rep I-III; n = 3).

| Vector | Transformation efficiency [%] | | | Transformation efficiency (mean ± SD) [%] |
|---|---|---|---|---|
| | Rep I | Rep II | Rep III | |
| FsPDS-KO1 | 11.20 | 17.29 | 10.28 | 12.92 ± 3.81 |
| FsPDS-KO2 | 8.43 | 6.53 | 14.34 | 9.79 ± 4.05 |

**Table S6: Deletions detected within ±10 bp of the cut site for *FsPDS*-targeting crRNAs in the wildtype control.** Columns indicate the position in the reference sequence, deletion length, deleted bases, number of reads with the deletion, and total reads covering the related reference position in the respective mapping.

| crRNA | Reference position | Length (bp) | Reference | Count | Total reads |
|---|---|---|---|---|---|
| 3_1 | 174 | 1 | G | 4 | 64006 |
| 3_1 | 177-178 | 2 | CA | 3 | 64006 |
| 3_1 | 180 | 1 | A | 1 | 64006 |
| 3_1 | 192 | 1 | T | 5 | 64006 |
| 3_2 | 192 | 1 | T | 5 | 63899 |
| 3_2 | 199 | 1 | C | 4 | 63899 |
| 3_2 | 200 | 1 | A | 1 | 63899 |
| 3_2 | 206 | 1 | T | 1 | 63899 |
| 3_2 | 208 | 1 | T | 3 | 63899 |
| 5_1/5_2 | 147 | 1 | T | 1 | 81207 |
| 5_1/5_2 | 148 | 1 | A | 2 | 81207 |
| 5_1/5_2 | 151 | 1 | C | 1 | 81207 |
| 5_1/5_2 | 153 | 1 | T | 1 | 81207 |
| 5_1/5_2 | 154 | 1 | G | 3 | 81207 |
| 5_1/5_2 | 156 | 1 | A | 1 | 81207 |
| 5_1/5_2 | 160 | 1 | T | 1 | 81207 |

**Table S7: Level-0 (L0) modules used for cloning of L1 promoter-reporter constructs.** L0 modules are derived from the MoClo Plant Parts Kit (Addgene #1000000047) and vector backbones from the MoClo Toolkit (Addgene #1000000044). Sequences for additional L0 modules are derived from vectors obtained from H. Puchta, KIT, Karlsruhe, Germany, or DNA Cloning Service (DCS), Hamburg, Germany.

| Name | Module | Destination | Source |
|---|---|---|---|
| pICH47811 | L1 Backbone vector | All L1 *mEGFP* vectors | Toolkit |
| pICH47802 | L1 Backbone vector | *Pc*Ubi::*turboRF*P | Toolkit |
| pICH41295 | L0 backbone vector | All L0 promoter vectors | Toolkit |
| pICH41308 | L0 backbone vector | L0.C13_*mEGFP* | Toolkit |
| pICH41276 | L0 backbone vector | L0.T12_*At*HSP18.2 | Toolkit |
| pICH51266 | CaMV35S+TMVΩ promoter | 35S::*mEGFP* | Plant Parts Kit |
| pICH87633 | *Atu*NOS promoter | *Atu*NOS::*mEGFP* | Plant Parts Kit |
| pICH88103 | *Atu*OCS promoter | *Atu*OCS::*mEGFP* | Plant Parts Kit |
| L0.P13_PcUbi | *Pc*UBI promoter | *Pc*UBI::*turboRFP* | KIT |
| L0.T12_pea3A | *Pea*3A terminator | *Pc*UBI::*turboRFP* | KIT |
| L0.P13_AtUbq10 | *At*UBQ10 promoter | *At*UBQ10::*mEGFP* | DCS |
| L0.P13_HaUbi | *Ha*UBI promoter | *Ha*UBI::*mEGFP* | DCS |
| L0.T12_AtHSP18.2 | *At*HSP18.2 terminator | All L1 *mEGFP* vectors | DCS |
| L0.C13_mEGFP | *mEGFP* CDS | All L1 *mEGFP* vectors | DCS |

**Table S8: Modifications of Level-0 modules for MoClo compatibility.**

| Module | Position (bp) | Base exchange | RE site | Mutation type |
|---|---|---|---|---|
| *mEGFP* CDS | 816 | G → A | *Bpi*I | silent |
| *At*UBQ10 promoter | 1279 | C → G | *Bpi*I | intron |
| *Ha*Ubi promoter | 167 | C → A | *Bsa*I | promoter |

**Table S9: Primers used in this study.**

| Identifier | Primer sequence (5’ → 3’) | Application |
|---|---|---|
| 3418 | gtagaagactgggagtttccttacattctgagc | Cloning of L0.P13_AtUbq10 |
| 3419 | atggaagactgcctcttcgatctaagattaa | Cloning of L0.P13_AtUbq10 |
| 3420 | gtagaagactcgaggattttctgggtttgatcg | Cloning of L0.P13_AtUbq10 |
| 3421 | gtagaagacgtcattctgttaatcagaaaaact | Cloning of L0.P13_AtUbq10 |
| 3531 | tctaagctctccaaagaccccaacgagaagagg | Cloning of L0.C13_mEGFP |
| 3532 | tcgttggggtctttggagagcttagattg | Cloning of L0.C13_mEGFP |
| 3552 | gtagaagacgcggaggacaaaacttaagaaagtcacc | Cloning of L0.P13_HaUbi |
| 3553 | gtagaagacgtggtagtagaccaacaaatggtatc | Cloning of L0.P13_HaUbi |
| 3556 | gtagaagactgaatgggcaagggcgaggaac | Cloning of L0.C13_mEGFP |
| 3557 | gtagaagactgaagctcacttgtagagttcatccatgccatgc | Cloning of L0.C13_mEGFP |
| 3558 | gtagaagactggctttgtcaaataaatagcttgtgtgc | Cloning of L0.T12_AtHSP18.2 |
| 3559 | gatgaagactgagcgagctccttatctttaatcatattcc | Cloning of L0.T12_AtHSP18.2 |
| 3560 | gtagaagactgggagaaaaattacggatatgaatataggcatatc | Cloning of L0.P13_PcUbi |
| 3561 | gtagaagactgcattgctgcacatacataacatatcaag | Cloning of L0.P13_PcUbi |
| 3562 | gtagaagactggcttcaggcctcccagctttc | Cloning of L0.T12_pea3A |
| 3563 | gtgaagactgagcgaagcctatactgtacttaacttgattgc | Cloning of L0.T12_pea3A |
| 3582 | gtagaagacggtaccacttacgtaacattaatattccatccg | Cloning of L0.P13_HaUbi |
| 3583 | gtagaagacggcattaagacgaagcgaaacaggaagagataag | Cloning of L0.P13_HaUbi |
| 3590 | gtagaagacttggaggaaccgcaacgttgaagg | Cloning of L0.P13_AtuNOS |
| 3591 | gtagaagacgccattgactctaattggataccgag | Cloning of L0.P13_AtuNOS |
| 3514 | ggttgcaaacatgagctggcac | Sequencing *FsPDS* Exon 1-3 |
| 3792 | ggacaaaaacagagcagcatg | Sequencing *FsPDS* Exon 1-3 |
| 3793 | tgttccatctcttggttaggg | Sequencing *FsPDS* Exon 4+5 |
| 3794 | cagggagagtgagggcag | Sequencing *FsPDS* Exon 4+5 |
| ttCas12a_for | gttcaccaactgctacagcc | ttCas12a amplification |
| ttCas12a_rev | gttcttgttgagggtgttgc | ttCas12a amplification |
| Ex3_for | agttaaggtactcttgggagatc | Exon 3 amplification |
| Ex3_rev | gattttctcattgcatcctgtacc | Exon 3 amplification |
| Ex5_for | ggcaagttggagcttacc | Exon 5 amplification |
| Ex5_rev | gatgaaatttgtccaccttctacg | Exon 5 amplification |

### Table S10: Edited reference sequences of *FsPDS* exons 3 and 5 used for alignment of amplicon sequencing data.

| Exon | *FsPDS* reference sequence |
|---|---|
| Exon 3 | AGTTAAGGTACTCTTGGGAGATCTAATAGTACTAGCTTAATGGATCATTACTTTTTTTTTTTTTTTTTCYT<br>TTTTCTTTTCATTTTCYTTTAATTGTTGCACTATASAGTATAATTTTACAAATTATTGCAGGTTTGGCTGG<br>TTTATCAACTGCCAAGTATTTGGCAGATGCTGGTCACAAACCTATACTATTGGAGTCAAGAGATGTTCTAG<br>GAGGAAAGGTTTTCATGCTGCTCTGTTTTTGTCCCACTTTTTCAATCAGCTGACACGAAACAAGTGCTTGT<br>TTTTCATTTTTGCCTTGCTTGCCACTTTCAAAAACTAGAATTTTTTTTTTTTGGATTGGTAATAAAAATTT<br>CATCAAATGATGAGAGAAATATACATGGGAAACCATTGGTACAGGATGCAATGAGAAAATC |
| Exon 5 | CATTACAAAATTAGGTTGTTTCCTTATTTGATTTTTTTTTGTTGGCAAGTTGGAGCTTACCCAAATGTGCA<br>GAATCTGTTTGGRGAACTTGGTATTAATGATCGGTTGCAATGGAAGGAACATTCTATGATATTTGCGATGC<br>CGAATAAGCCTGGAGAGTTCAGCCGATTTGATTTTCCYGAAGTTCTTCCTGCACCATTAAATGGTATTATA<br>AGAGTAATCACTAGTGCGGCCGCCTGCAGGTCGACCATATGGGAGAGCTCCCAACGCGTTGGATGCATAGC<br>TTGAGTATTCTATAGTGTCACCTAAATAGCTTGGCGTAATC |
