## Supplemental Figures for "First report of genetic transformation and CRISPR/Cas12a-mediated gene editing of European beech (*Fagus sylvatica* L.) employing a transient protoplast system"

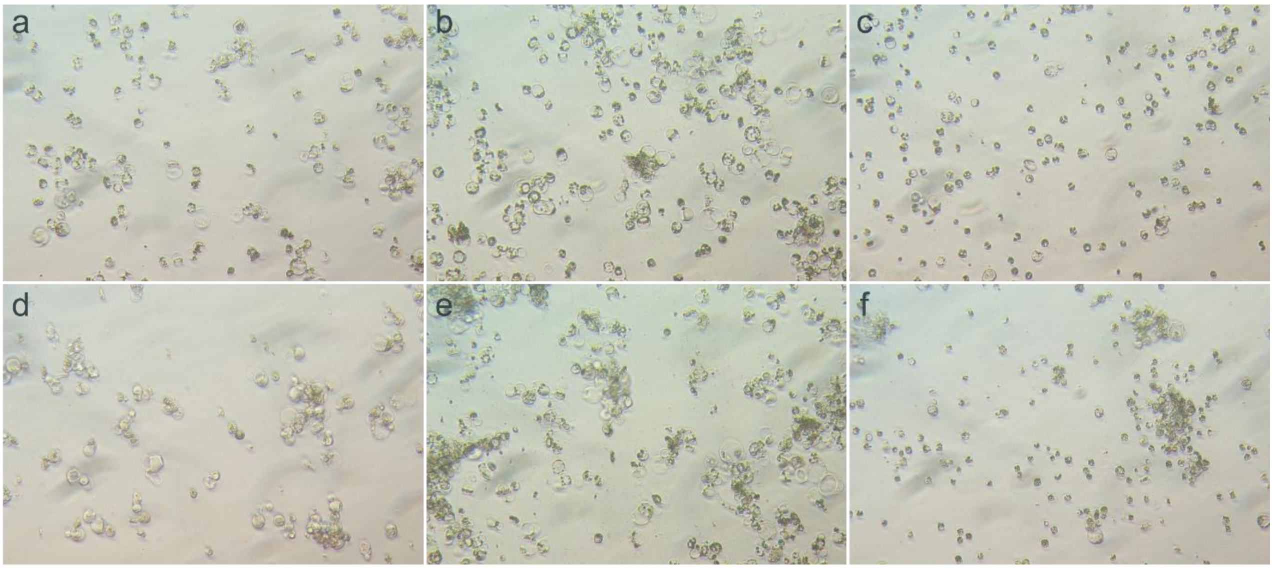


**Figure S1: Beech protoplast viability after 5 (a-c) and 9 days (d-f) in different media**: regeneration medium (a, d), regeneration medium + antibiotics (b, e), regeneration medium + antibiotics + natamycin (c, f). Representative images illustrate effects on survival and morphology of two individual isolations.

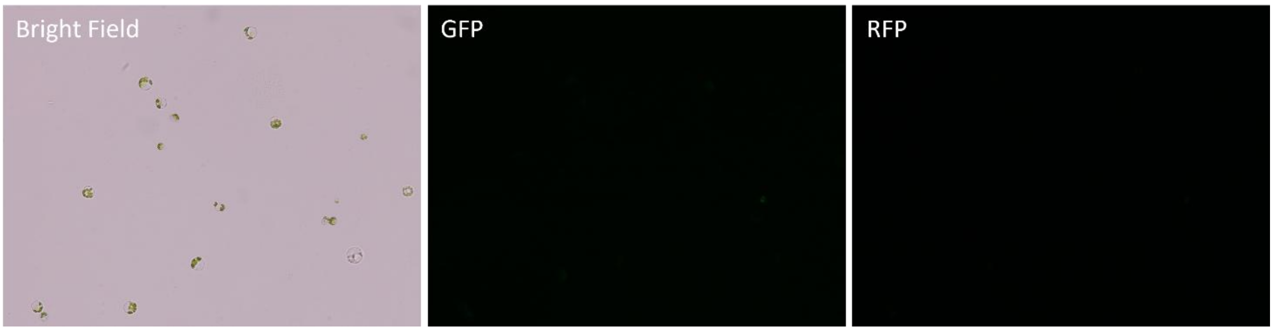


**Figure S2: Negative control of protoplast transfection for promoter-reporter validation**. Bright-field, GFP, and RFP channels of protoplasts transfected without plasmid DNA, imaged under the same conditions as experimental samples. No fluorescence signal was detected in the GFP or RFP channels, confirming the absence of autofluorescence or nonspecific signal.

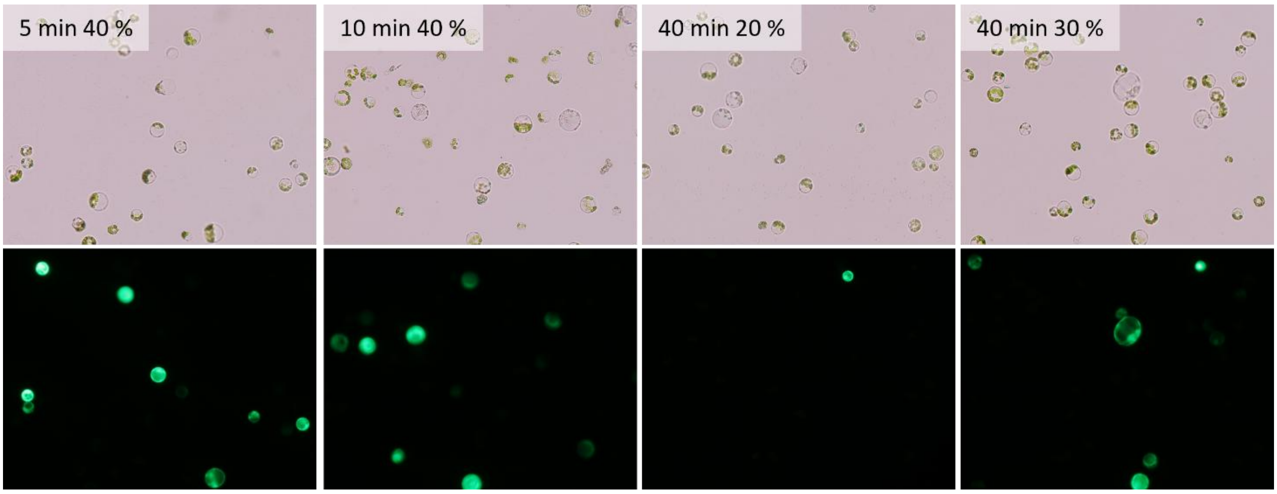


**Figure S3: Protoplast morphology and mEGFP expression of *F. sylvatica* protoplasts 24 h post-transformation using PEG$_{1500}$.** The panels illustrate the acute effect of varying PEG$_{1500}$ concentration and incubation time on protoplast morphology and subsequent transformation success. Upper panels (bright-field microscopy): protoplast morphology was assessed to evaluate acute PEG-induced cytotoxicity. High vitality was characterized by the maintenance of a spherical shape. Conversely, signs of toxicity are increased morphological damage, including irregularly shaped protoplasts, shrunken cells, and cell debris. The incubation conditions are indicated above each panel (min and PEG$_{1500}$ concentration [%]). The lower panels (green fluorescence) confirmed the success of transformation (e.g., mEGFP expression) under these conditions.

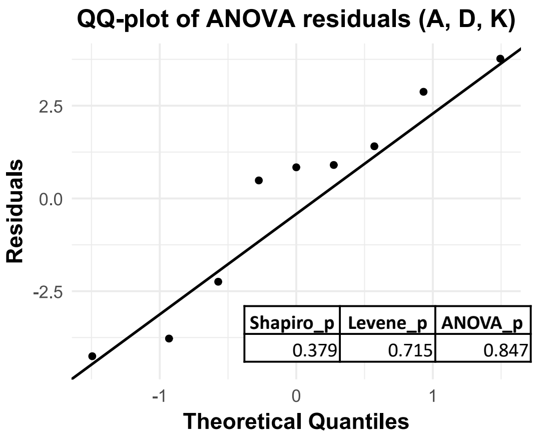


**Figure S4: QQ-plot of ANOVA residuals for protoplast transformation efficiencies (parameters A, D, and K).** The QQ-plot confirms the approximate normal distribution of the residuals, fulfilling a key assumption for the one-way ANOVA. Statistical test results supporting the model's assumptions and the main comparison are presented in the inset table: Shapiro_p (0.379) ≥ 0.05 indicates the residuals are normally distributed. Levene_p (0.715) ≥ 0.05 indicates homogeneity of variances. ANOVA_p (0.847) ≥ 0.05 indicates no significant difference in mean transformation efficiency between parameter sets A, D, and K (n = 3 biological replicates per set).

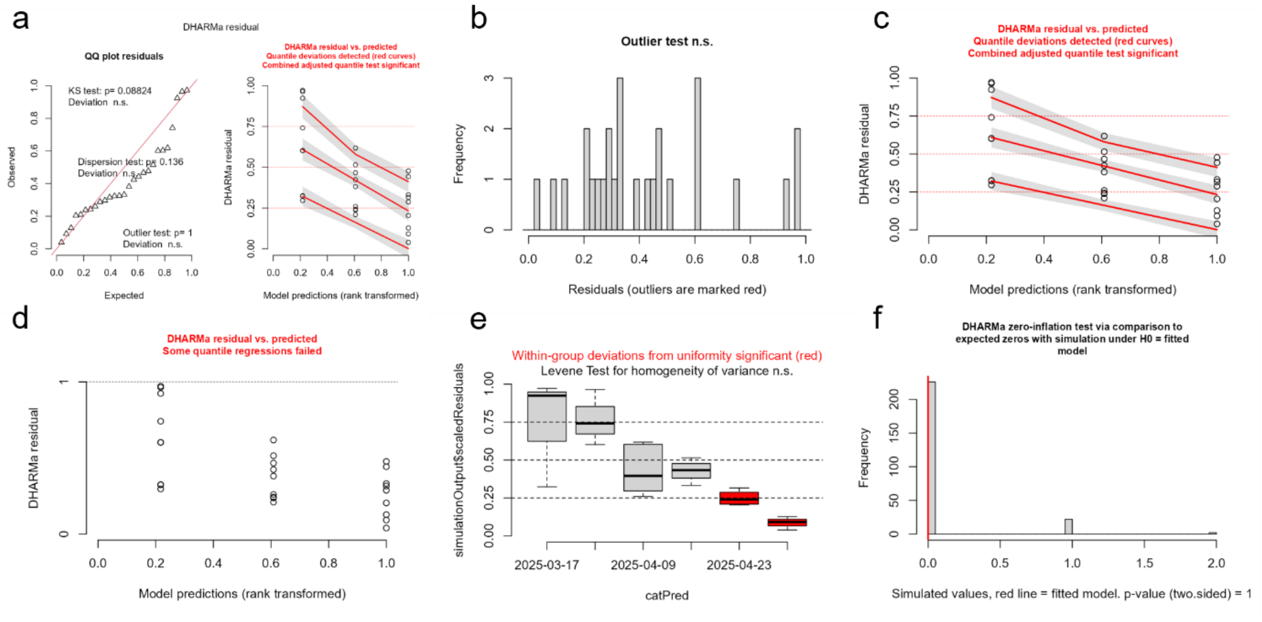


**Figure S5: Diagnostic plots of GLMM for comparison of transformation parameters over time.** The plots show the model diagnostics (a-f) for the Generalized Linear Mixed Model (GLMM). (a) The QQ-plot indicates a significant deviation from the theoretical distribution, suggesting a lack of fit. (b) The outlier test shows no significant outliers. (c) The quantile residuals plot reveals significant deviations, indicating that the model's assumptions about variance are not fully met. (d) Some quantile regressions failed, further highlighting model limitations. (e) The within-group deviations from uniformity are significant, a potential issue with the nested data structure. (f) The zero-inflation test suggests the model is a good fit for the number of zeros.

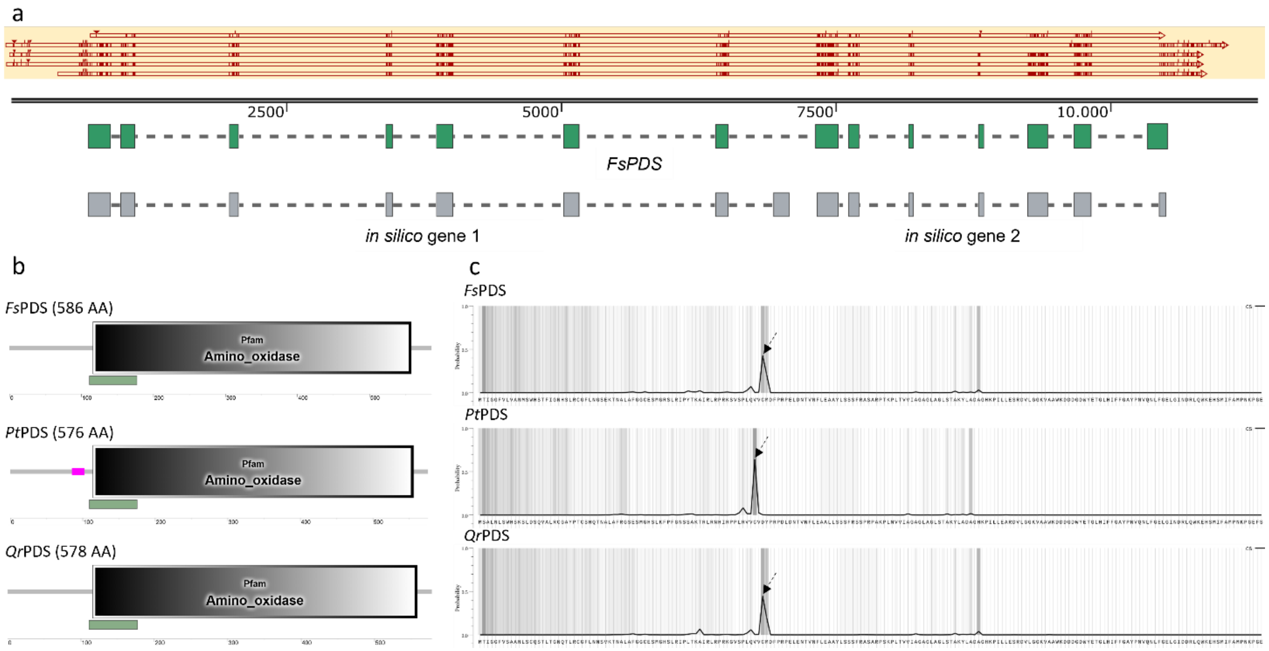


**Figure S6: Annotation of *F. sylvatica PHYTOENE DESATURASE* (*FsPDS*).** (a) Exon–intron boundaries were determined by sequence alignments with *PDS* sequences (shown from top to bottom): CDS from *Populus trichocarpa* (*PtPDS*; Potri.014G148700), mRNA from *Quercus suber* (*QsPDS*; NCBI RefSeq: XM_065777133.1), *Quercus robur* (*QrPDS*; NCBI RefSeq: XM_050436637.1), *Castanea sativa* (*CsPDS*; NCBI RefSeq: XM_075797933.1), and *Quercus lobata* (*QlPDS*; NCBI RefSeq: XM_031066959.1). Additionally, gene models were predicted *in silico* with FGENESH (Softberry). The final gene structure comprises 14 exons. (b) The translated protein sequence was verified by assessing the completeness of PDS-type functional domains (Flavin-containing amine oxidoreductase (Amino-oxidase; PF01593), NAD(P)-binding Rossmann-like domain (PF13450; green bar), low complexity region (pink bar)) using SMART. (c) The presence of a chloroplast transit peptide (arrow) within the PDS proteins was predicted with TargetP-2.0.

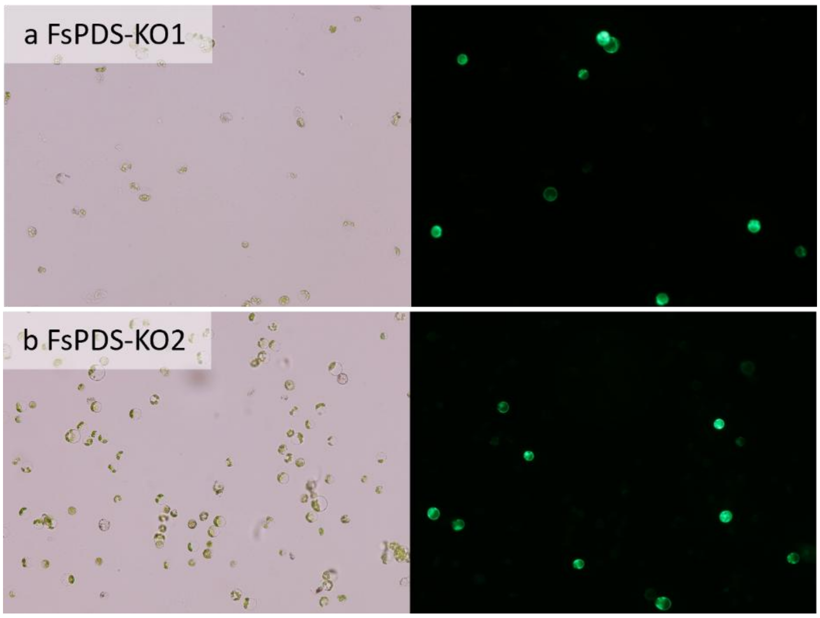


**Figure S7: *F. sylvatica* protoplasts after transformation with the CRISPR/Cas12a vectors.** Representative images of protoplasts 48 hours after transformation with FsPDS-KO1 (KO1, a) or FsPDS-KO2 (KO2, b). The figure shows the bright-field image (left) and the GFP fluorescence image (right). Green fluorescence confirmed the successful DNA transfer.

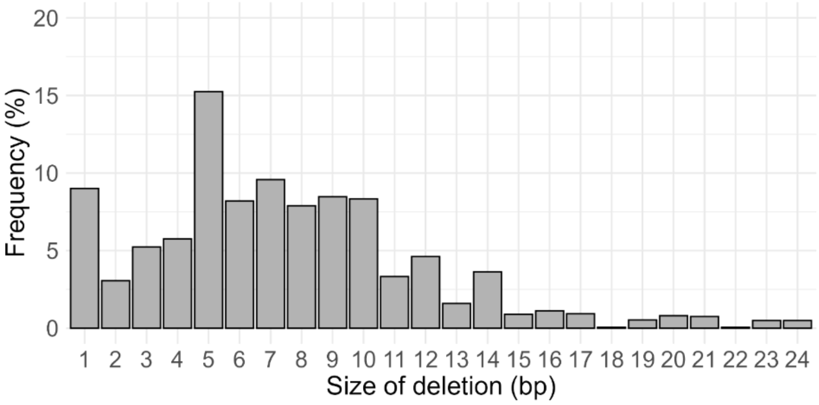


**Figure S8: Cumulative distribution of deletion sizes induced by tt*Lb*Cas12a across four crRNAs targeting *FsPDS* in *F. sylvatica* protoplasts.** Deletion sizes were aggregated across all 4 crRNA–target combinations. Values represent the summed, normalized fraction of reads for each deletion size. The distribution shows a strong enrichment of small to mid-sized deletions, with the majority of deletion events concentrated between approximately 4 and 10 bp.
