## Supplementary material for "First report of genetic transformation and CRISPR/Cas12a-mediated gene editing of European beech (*Fagus sylvatica* L.) employing a transient protoplast system": Data S3 - Statistical data: ANOVA_ADF.R

```
# =========================================
# load packages
# =========================================
library(tidyverse)
library(writexl)
library(car)       #Levene-Test
library(ggplot2)

# =========================================
# 1. Read and prepare data
# =========================================
df <- Transformation_effciencys_statistics_A_D_F %>%
  mutate(Efficiency_Replica = (sum_GFP / sum_Cells) * 100)

# =========================================
# 2. Aggregate over technical replicates (mean per parameter × date)
# =========================================
agg <- df %>%
  group_by(Parameter, Date) %>%
  summarise(
    mean_eff = mean(Efficiency_Replica, na.rm = TRUE),
    .groups = "drop"
  )

# =========================================
# 3. ANOVA über Parameter
# =========================================
anova_res <- aov(mean_eff ~ Parameter, data = agg)
anova_p   <- summary(anova_res)[[1]][["Pr(>F)"]][1]


# =========================================
# 4. Residuen für Tests
# =========================================
resid_vals <- residuals(anova_res)

# Shapiro-Wilk-Test auf Residuen
shapiro_res <- shapiro.test(resid_vals)
shapiro_df <- tibble(
  Shapiro_W = shapiro_res$statistic,
  Shapiro_p = shapiro_res$p.value
)

# Levene-Test auf Gruppen
levene_raw <- leveneTest(mean_eff ~ Parameter, data = agg)
levene_df <- tibble(
  Levene_F = levene_raw$`F value`,
  df1 = levene_raw$Df[1],
  df2 = levene_raw$Df[2],
  Levene_p = levene_raw$`Pr(>F)`[1]
)

# =========================================
# 5. QQ-Plot of residues
# =========================================
qq_plot <- ggplot(data.frame(resid = resid_vals), aes(sample = resid)) +
  stat_qq(size = 2) +                 # etwas größere Punkte
  stat_qq_line(size = 1) +            # etwas dickere Linie
  labs(
    title = "QQ-plot of ANOVA residuals (A, D, F)",
    x = "Theoretical Quantiles",
    y = "Residuals"
  ) +
  theme_minimal(base_size = 16) +     # Grundschriftgröße größer
  theme(
    plot.title = element_text(size = 20, face = "bold", hjust = 0.5),
    axis.title = element_text(size = 18, face = "bold"),
    axis.text = element_text(size = 14)
  )

# save QQ-Plot as .PNG
png("QQPlot_Residuen_A_D_F.png", width = 900, height = 700, res = 150)
print(qq_plot)
dev.off()

# =========================================
# 6. Post-hoc Tukey (only if ANOVA is significant; p < 0.05)
# =========================================
if (anova_p < 0.05) {
  tukey_raw <- TukeyHSD(anova_res, "Parameter")$Parameter
  posthoc_df <- as.data.frame(tukey_raw) %>%
    rownames_to_column("contrast") %>%
    rename(estimate = diff, lwr = lwr, upr = upr, p.value = `p adj`) %>%
    mutate(
      Signifikanz = case_when(
        p.value < 0.001 ~ "***",
        p.value < 0.01  ~ "**",
        p.value < 0.05  ~ "*",
        TRUE            ~ ""
      )
    )
} else {
  posthoc_df <- tibble(contrast = NA, estimate = NA, lwr = NA, upr = NA,
                       p.value = NA, Signifikanz = NA)
}

# =========================================
# 7. export results
# =========================================
write_xlsx(list(
  "Replica" = agg,
  "ANOVA" = tibble(ANOVA_p = anova_p),
  "Posthoc_TukeyHSD" = posthoc_df,
  "Shapiro_Residuals" = shapiro_df,
  "Levene_Residuals" = levene_df
), "ANOVA_parameter.sets_ADF.xlsx")

# princt for control
print(tibble(ANOVA_p = anova_p))
print(posthoc_df)
print(shapiro_df)
print(levene_df)
```
