## Supplementary material for "First report of genetic transformation and CRISPR/Cas12a-mediated gene editing of European beech (*Fagus sylvatica* L.) employing a transient protoplast system": Data S3 - Statistical data: GLMM_ADF+Plot.R

```
# =========================================
# 📦 Load required packages
# =========================================
library(tidyverse)
library(lme4)
library(lmerTest) 
library(emmeans)
library(writexl)
library(DHARMa)
library(ragg)

# =========================================
# 1. Data preparation
# =========================================
# Replace with your dataset
df_raw <- Transformation_effciencys_statistics_A_D_F

df <- df_raw %>%
  mutate(
    TPP = as.factor(Date),
    Parameter = as.factor(Parameter),
    obs_id = 1:n()  # unique observation ID for overdispersion correction
  ) %>%
  droplevels()

# =========================================
# 2. GLMM analysis with DHARMa diagnostics
# =========================================
# Output directories
dir.create("GLMM_Diagnostics", showWarnings = FALSE, recursive = TRUE)

# Initial GLMM
glmm_initial <- glmer(
  cbind(sum_GFP, sum_Cells - sum_GFP) ~ Parameter + (1 | TPP),
  data = df,
  family = binomial
)

# Test for overdispersion
sim_out <- simulateResiduals(glmm_initial, plot = FALSE)
disp_test <- testDispersion(sim_out)

# Final model (add obs_id random effect if overdispersion detected)
glmm_final <- if (disp_test$p.value < 0.05) {
  message("Overdispersion detected. Adding observation-level random effect.")
  glmer(
    cbind(sum_GFP, sum_Cells - sum_GFP) ~ Parameter + (1 | TPP) + (1 | obs_id),
    data = df,
    family = binomial
  )
} else {
  message("No significant overdispersion detected. Using initial model.")
  glmm_initial
}

# DHARMa diagnostics
sim_out_final <- simulateResiduals(glmm_final, plot = FALSE)
ragg::agg_png("GLMM_Diagnostics/dharma_all_diagnostics.png",
              width = 1600, height = 1200, units = "px", res = 150)
plot(sim_out_final, testOutliers = TRUE, testDispersion = TRUE, testZeroInflation = TRUE)
dev.off()

# Summarize DHARMa tests
dharma_tests <- tibble(
  Test = c("Uniformity", "Dispersion", "Outliers", "ZeroInflation"),
  p_value = c(
    testUniformity(sim_out_final)$p.value,
    testDispersion(sim_out_final)$p.value,
    testOutliers(sim_out_final, plot = FALSE)$p.value,
    testZeroInflation(sim_out_final, plot = FALSE)$p.value
  )
)

# =========================================
# 3. Model evaluation & EMMs
# =========================================
# Likelihood-ratio test
glmm_null <- update(glmm_final, . ~ . - Parameter)
anova_glmm <- anova(glmm_null, glmm_final)

# Estimated marginal means (EMMs)
emm_glmm <- emmeans(glmm_final, ~ Parameter, type = "response")

# Prepare EMMs for plotting
emm_df <- summary(emm_glmm) %>%
  as.data.frame() %>%
  mutate(
    response_percent = prob * 100,
    CI_lower = asymp.LCL * 100,
    CI_upper = asymp.UCL * 100
  )

# Pairwise post-hoc comparisons
pairwise_glmm <- pairs(emm_glmm, adjust = "tukey") %>% as.data.frame()

# Export results
write_xlsx(list(
  "ANOVA_LRT" = as.data.frame(anova_glmm),
  "Posthoc_Tukey" = pairwise_glmm,
  "EMM_percent" = emm_df,
  "DHARMa_Tests" = dharma_tests
), "Supplemental_GLMM_Results.xlsx")

# =========================================
# 4. Plot raw data & EMMs
# =========================================
dir.create("GLMM_Plots", showWarnings = FALSE)

tpp_colors <- c(
  "2025-03-17" = "#2A788E",
  "2025-03-27" = "#7AD151",
  "2025-04-09" = "#FDE725",
  "2025-04-16" = "#2A788E",
  "2025-04-23" = "#7AD151",
  "2025-04-28" = "#FDE725"
)

p <- ggplot(df, aes(x = Parameter, y = (sum_GFP/sum_Cells)*100)) + 
  geom_violin(fill = "white", trim = FALSE, alpha = 0.8, color = "black", linewidth = 0.6) +
  geom_jitter(aes(fill = TPP, shape = TPP), width = 0.15, size = 5, stroke = 0.2, color = "black") +
  geom_pointrange(data = emm_df, 
                  aes(x = Parameter, y = response_percent, ymin = CI_lower, ymax = CI_upper),
                  color = "black", size = 1.5, linewidth = 1, width = 0.5) +
  scale_fill_manual(values = tpp_colors) +
  scale_shape_manual(values = c(21, 21, 21, 22, 22, 22)) +
  labs(y = "Transformation efficiency (%)", x = NULL, fill = "Date", shape = "Date") +
  ylim(0, 25) +
  theme_minimal(base_size = 16) +
  theme(
    text = element_text(family = "Arial"),
    legend.position = "right",
    legend.title = element_text(size = 18, face = "bold"),
    legend.text  = element_text(size = 20),
    axis.text.x = element_text(size = 24, angle = 45, hjust = 1),
    axis.text.y = element_text(size = 24),
    axis.title.y = element_text(size = 24, face = "bold"),
    plot.margin = margin(10, 10, 10, 10)
  )

ragg::agg_png("GLMM_Plots/GLMM_Plot.png", width = 1200, height = 800, units = "px", res = 96)
print(p)
dev.off()
```
